# Targeted inhibition of colorectal carcinoma using a designed CEA-binding protein to deliver TCF/LEF transcription factor decoy DNA and p53 protein

**DOI:** 10.1101/2025.06.09.658751

**Authors:** Wen Wang, Xuan Sun, Geng Wu

**Author notes:** To whom correspondence may be addressed (G.W.).

## Abstract

Colorectal carcinoma (CRC) is characterized by mutations in Wnt signaling pathway components and the p53 protein, and anti□PD-1/anti□PD-L1 immunotherapy has shown limited efficacy in microsatellite stable CRC patients. In this work, CEABP1, a de novo-designed binding protein for the CRC marker carcinoembryonic antigen (CEA), and cell-penetrating peptide (CPP) were employed for targeted delivery in CRC cells. The consensus binding motifs for TCF/LEF and Max were concatenated, and Max DNA binding domain protein fused with CPP and CEABP1 was used to deliver this TCF/LEF transcription factor decoy (TFD) DNA specifically into CRC cells by recognizing CEA on the CRC cell surface to inhibit Wnt target gene transcription, leading to marked suppression of CRC cell proliferation and xenograft tumor growth. In addition, p28 was employed as a CPP to specifically deliver the p28-p53-CEABP1 protein into CRC cells, which significantly enhanced p53 inhibition of CRC cell proliferation and xenograft tumor growth. Furthermore, codelivery of the p14^ARF^ protein together with p53 increased its antitumor activity by prolonging its effective duration. These findings paved the way for the development of related biomacromolecular anticancer therapeutics.

## Introduction

Colorectal carcinoma (CRC) is the third most diagnosed cancer worldwide, with almost two million new cases and more than 903 thousand deaths each year.^1,2^ CRC is commonly categorized into nonhypermutated and hypermutated CRC. Nonhypermutated CRC is usually microsatellite stable (MSS) but chromosome instable (CIN), accounting for 84% of CRC cases. In contrast, hypermutated CRC, which accounts for 16% of CRC cases, is generally characterized by microsatellite instability (MSI), with mutations in DNA mismatch repair (dMMR)-related proteins.^3,4,5^ For hypermutated CRC with dMMR or high MSI, anti□PD-1/anti□PD-L1 immunotherapy has been proven to be effective. However, for the MSS type of CRC, which constitutes the majority of CRC cases, the efficacy of immune checkpoint inhibitors is limited.^6^

The occurrence of CRC is generally the result of multiple genetic mutations. First, CRC is highly correlated with mutations in proteins of the Wnt signaling pathway, which occur in more than 94% of CRC cases, with biallelic inactivating mutations of *APC*, for example.^3^ These mutations result in abnormally elevated levels of β-catenin in the cytoplasm, which translocates into the nucleus and interacts with the transcription factor T-cell factor (TCF)/lymphoid enhancer-binding factor (LEF), promoting the transcription of target genes such as *cyclin D1* and *c-myc* and leading to abnormal cell proliferation.^7,8,9^ Mutations of *APC* typically occur in benign adenoma stages.^10,11^ Second, mutations in the tumor suppressor p53 frequently occur in CRC, which usually happens in late adenoma or carcinoma stages.^11,12^ Encoded by the *TP53* gene, the p53 protein is known as the “guardian of the genome”, and 50% of all cancer patients are found to have p53 mutations.^13,14^ In 60% of nonhypermutated CRC cases, mutations in p53 were detected. p53 functions mainly as a transcription factor, promoting the expression of p21^Cip^^1^ (encoded by *cdkn1a*) to inhibit cyclin-CDK and cause cell cycle arrest, as well as stimulating the expression of *bax*, *noxa*, and *puma* to lead to cancer cell apoptosis. In addition, p53 functions independently of transcription.^14^ The intracellular protein level of p53 is maintained at a low level by several E3 ubiquitin ligases, including Mdm2, ARF-BP1/Mule, COP1, and Pirh2, which polyubiquitinate p53 for degradation by the 26S proteasome.^15–19^ Another tumor suppressor, p14^ARF^, interacts with both Mdm2 and ARF-BP1/Mule to counteract their polyubiquitination of p53 so that p53 is prevented from degrading.^20–24^ Third, other genes, including *kras*, *pik3ca*, *smad4*, etc., are also frequently mutated in CRC.

Although mutations in proteins often lead to excessive activation of the Wnt signaling pathway in colorectal cancer, there are currently no effective means to antagonize overactivated Wnt signaling.^25^ The potential of nucleic acids as a therapeutic means to treat diseases is continuously being explored. The current nucleic acid-based therapeutics include antisense oligonucleotides (ASOs), aptamers, small interfering RNAs (siRNAs), CRISPR/Cas9-based gene editing, and transcription factor decoys.^26,27,28^ In transcription factor decoy (TFD) technology, a short piece of double-stranded DNA, which represents the consensus binding site for the target transcription factor, is added to cells, where it competitively interacts with endogenous transcription factor proteins and prohibits the transcription of target genes.^29,30,31^ However, owing to the difficulty of cancer cell-specific delivery of DNA, TFD has not yet been widely applied in anticancer therapeutics.

In current cancer treatment, intervention measures, such as small molecule inhibitors, siRNAs, and PROTACs, exist only for oncoprotein mutations. On the other hand, for tumor suppressor protein mutations, there are no suitable drugs available. Antibodies are extracellular proteins that generally function outside cells, while anticancer therapeutics based on intracellular proteins have not yet been well developed. A promising strategy is to use cell-penetrating peptides (CPPs) or other means to deliver purified tumor suppressor proteins into tumor cells to inhibit cancer cell proliferation.^32^ Matsui *et al*. used polyarginines,^33,34^ whereas Choi *et al*. employed HIV-1 TAT as a CPP to deliver the p53 protein into cancer cells to restore p53 function;^35^ Tang *et al*. applied polyethylene glycol nanocapsules to facilitate p53 protein entry into cancer cells;^36^ and Chan *et al*. reported that engineered *Bacillus thuringiensis* Cry3Aa protein crystals could be used to deliver the p53 protein to sensitize triple-negative breast cancer cells to anti-PD-1 immunotherapy.^37,38^ However, these works, which mostly employ cell culture without thorough in vivo investigations, have more or less suffered from inefficient lysosomal escape after protein delivery and lack specific targeting for cancer cells. In addition, the p53 protein is not stable because of the actions of multiple E3 ubiquitin ligases, such as Mdm2. Therefore, anticancer therapeutics based on p53 protein delivery has not been widely developed.

In this work, TCF/LEF TFD DNA comprising a TCF/LEF-binding sequence concatenated with the Max-binding sequence was delivered into CRC cells via purified human Max-binding domain protein fused with a CPP to competitively interact with endogenous TCF/LEF in CRC cells and suppress Wnt signaling. Moreover, an artificial binding protein for the CRC marker CEABP1, carcinoembryonic antigen-related cell adhesion molecule 5 (CEACAM5, abbreviated as CEA hereafter), was designed de novo and fused to the CPP-Max binding domain protein to specifically deliver TCF/LEF TFD DNA into CRC cells to repress overactivated Wnt signaling and suppress CRC cell proliferation. Furthermore, CEABP1 was fused to p53 tagged with p28, a CPP derived from *Pseudomonas aeruginosa*,^39,40,41^ to specifically deliver purified p53 protein into CRC cells through endocytosis to stimulate the expression of p53 target genes, promote cell cycle arrest and apoptosis, and prevent CRC cell growth. In addition, the purified N-terminal domain of p14^ARF^ was delivered by CPP simultaneously with p53 to prolong its lifetime and increase its effective duration. These studies will contribute to the further development of related biomacromolecular anticancer therapeutics based on TFD DNA and purified intracellular tumor suppressor proteins.

## Results

### Delivery of TCF/LEF TFD DNA by the CPP-Max protein suppressed CRC cell proliferation and xenograft tumor growth

In 92% of nonhypermutated and 97% of hypermutated CRC cases, mutations in proteins in the Wnt signaling pathway were detected. Some of these mutations occur upstream of Wnt signaling, such as RNF43/ZNRF3; some exist midstream, such as APC and Axin; and some occur downstream, such as β-catenin or TCF4/TCF3.^3,5^ If we intervene upstream, overactivation of Wnt signaling due to mutations in midstream or downstream components would still not be inhibited. Therefore, we chose to intervene at the most downstream stage of Wnt signaling, which involves complex formation between the transcription factors TCF/LEF and their target gene promoters.

One of the difficulties in nucleic acid therapeutics is how to deliver nucleic acids into cells. Instead of using lipid nanoparticles, adenovirus-associated viruses, or dendrimers, our approach was to synthesize a short piece (36 base pairs, bp) of double-stranded DNA in which the binding sequence of TCF/LEF and that of a DNA-binding protein were concatenated. Next, the corresponding DNA-binding protein fused with a cell-penetrating peptide (CPP) was purified and employed to deliver the DNA into CRC cells to function as a TFD to competitively block endogenous TCF/LEF proteins from accessing the promoters of Wnt signaling-responsive genes (Figure 1A). Various DNA-binding proteins, including TetR, *Streptomyces coelicolor* ScoMcrA-SBD,^42^ and the human Max DNA-binding domain (abbreviated as Max hereafter), and different CPPs, such as TAT, p28, penetratin (pene), and pep1 were tested,^32^ and Max fused with pep1 and a nuclear localization sequence (abbreviated as pep1-Max hereafter) exhibited the best result in terms of DNA delivery and suppression of CRC cell proliferation; thus, it was selected for subsequent studies. Max dimers preferentially bind to promoter regions containing enhancer box (E-Box) sequences with a 5□-CACGTG-3□ consensus motif.^43^ TCF/LEF TFD DNA was synthesized in which the E-Box sequence was linked with the TCF/LEF consensus binding site 5□-AGATCAAAGG-3□, with phosphorothioation modification at the ends of the DNA to reduce degradation by exonucleases.^28^ After delivery via the purified pep1-Max protein, the fluorophore FAM-labeled TCF/LEF TFD DNA could be readily internalized into CRC cells and was localized in the nucleus, as indicated by the intense fluorescence signal in nearly all the cell nuclei (Figure 1B). Both the CCK-8 assay (Figure 1C) and the colony formation assay (Figures 1D and 1E) revealed that the delivered TCF/LEF TFD DNA efficiently suppressed CRC cell proliferation, whereas pep1-Max protein alone or DNA alone had no notable effects on cell growth.

**Figure 1.**
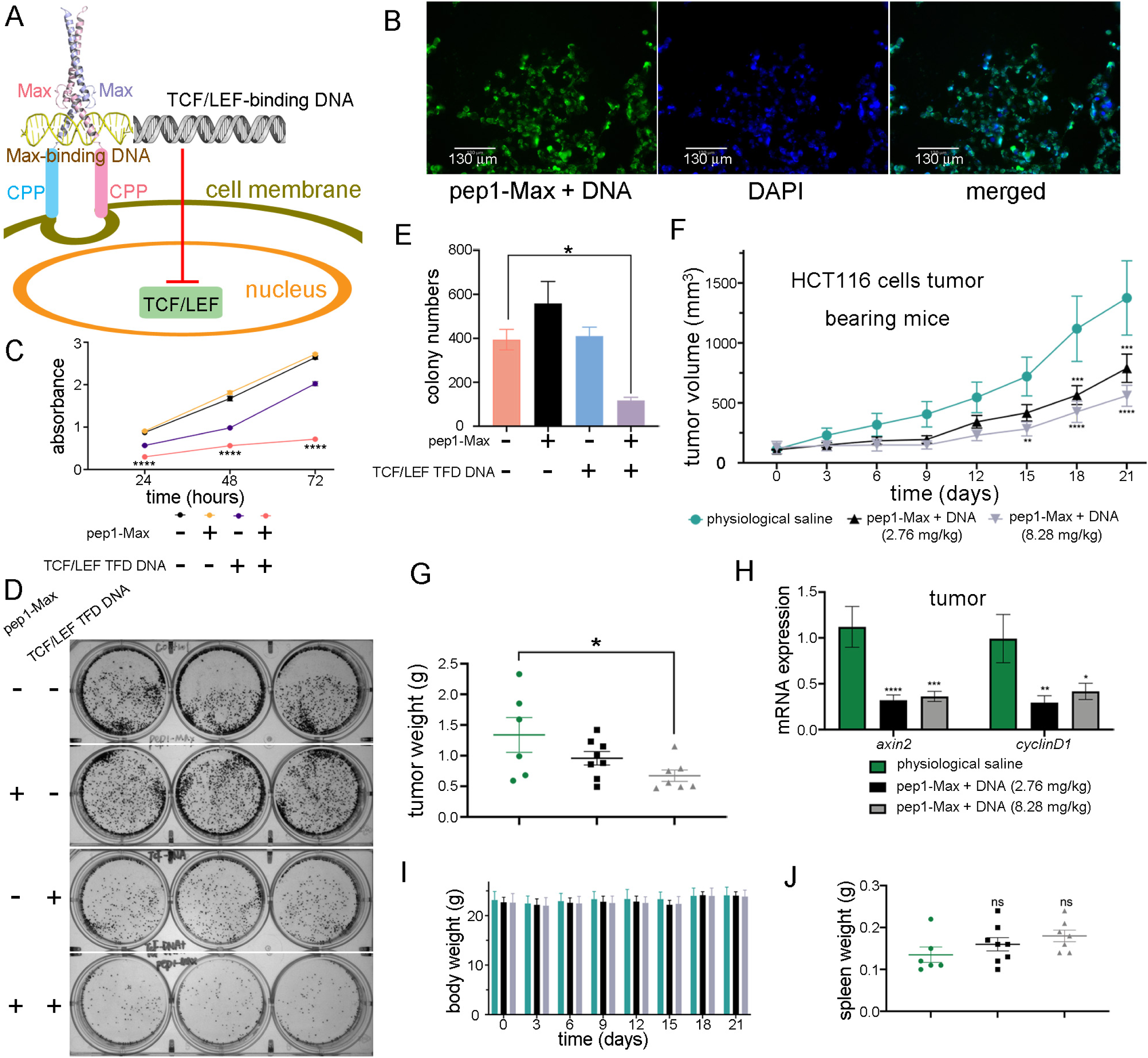
Delivery of TCF/LEF TFD DNA into CRC cells via the pep1-Max protein inhibited CRC cell proliferation and xenograft tumor growth in mice. (A) Delivery scheme of TCF/LEF TFD DNA. The TCF/LEF-binding DNA sequence was linked with the Max-binding DNA sequence as TCF/LEF TFD DNA. The human Max DNA-binding domain was fused with the cell-penetrating peptide pep1, and the fusion protein was employed to deliver the TCF/LEF TFD DNA into CRC cells, which competitively inhibited endogenous TCF/LEF from binding to its target gene promoters and prevented the transcription of Wnt signaling-responsive genes. (B) TCF/LEF TFD DNA could be efficiently delivered into LS174T CRC cells and was located in the nucleus. Fluorescence microscopy images of LS174T cells delivered with FAM-labeled TCF/LEF TFD DNA and purified pep1-Max protein are shown. Scale bars: 130 μm. (C) Delivery of TCF/LEF TFD DNA by the purified pep1-Max protein inhibited the proliferation of HCT-116 cells, as revealed by the CCK-8 assay. (D) Delivery of TCF/LEF TFD DNA by purified pep1-Max protein suppressed HCT116 cell proliferation in the colony formation assay. (E) Quantification of the colony formation assay results. (F) Treatment of HCT116 cell xenograft mice via tail vein injection of TCF/LEF TFD DNA and purified pep1-Max protein decelerated tumor growth. Tumor growth curves of HCT116 cell subcutaneous xenograft mice treated with physiological saline solution (n = 6), 2.76 mg/kg pep1-Max and TCF TFD DNA (molar ratio 3:1, n = 8), or 8.28 mg/kg pep1-Max and TCF TFD DNA (molar ratio 3:1, n = 7) are shown. The error bars represent the standard error of the mean (SEM), which was determined via two-way ANOVA. (G) Tumor weight at the experimental endpoint (day 21). The color scheme for different groups is the same as that in (F). (H) Treating mice with TCF/LEF TFD DNA and pep1-Max decreased *axin2* and *cyclin D1* mRNA expression in tumors, as revealed by qRT□PCR analysis of tumor samples. (I) Treatment of mice with TCF/LEF TFD DNA and pep1-Max protein did not affect the body weight of the mice during the experiment. The color scheme for the different groups is the same as that in (F). (J) Treatment of the mice with TCF/LEF TFD DNA and pep1-Max protein did not affect the spleen weight of the mice at the experimental endpoint (day 21). The color scheme for different groups is the same as that in (H).

The antitumor effect of TCF/LEF TFD DNA was further investigated in a xenograft tumor mouse model. HCT116 CRC cells were injected subcutaneously (s.c.) into immunodeficient BALB/c nude mice. After the tumors had grown to a certain size, TCF/LEF TFD DNA together with purified pep1-Max protein was systemically injected via the tail vein every 3 days for six consecutive treatments. Compared with the physiological saline control, a significant reduction in tumor volume was observed in the mice treated with TCF/LEF TFD DNA and pep1-Max protein (Figure 1F, Figure S1). At the end of the experiment, a significant decrease in tumor weight was observed between the group of mice receiving 8.28 mg/kg TCF/LEF TFD DNA and pep1-Max and the group receiving physiological saline (Figure 1G). Furthermore, the expression of the Wnt signaling target genes *axin2* and *cyclin D1* in tumor tissues was markedly downregulated by the treatment with TCF/LEF TFD DNA and pep1-Max compared with the physiological saline control (Figure 1H). Moreover, TCF/LEF TFD DNA treatment did not affect the body weight (Figure 1I) or spleen weight (Figure 1J) of the mice throughout the experiment, even at higher doses.

Taken together, these results showed that delivery of TCF/LEF TFD DNA via the purified pep1-Max protein effectively suppressed CRC cell proliferation and xenograft tumor growth.

### Targeting CEACAM5 (CEA)-expressing CRC cells with a designed CEA-binding protein enhanced the CRC-suppressing effect of TCF/LEF TFD DNA

A goal of continuous effort in cancer drug discovery is to specifically target tumor cells without harming normal cells. To achieve this goal, de novo protein design technology was applied to design a binding protein for the extracellular domain of a membrane CRC marker. Carcinoembryonic antigen-related cell adhesion molecule 5 (CEACAM5, abbreviated as CEA hereafter) is a well-known tumor marker of colorectal cancer that is highly expressed on the membranes of CRC cells.^44–45^ Alphafold3 was employed to predict the structure of the full-length CEA, which showed that it consists of seven extracellular immunoglobulin (IgG) domains, which is consistent with the present knowledge. These seven IgG domains are named the N, A1, B1, A2, B2, A3, and B3 domains, from distal to proximal to the cell membrane (Figure 2). As the N, A1, and B1 domains are relatively far from the cell membrane and the B3 domain is too close, the A3 and B2 domains of CEA were selected as the recognition targets of artificially designed proteins. RFDiffusion, which is based on diffusion models, was utilized to design the main-chain backbone scaffolds, and ProteinMPNN, which uses graph neural networks and a message-passing mechanism to integrate spatial and energy information, was employed to design the protein sequences (Figure 2). Using this scaffold-sequence two-step approach, binding proteins for the CEA-A3 domain were designed. One of them, named as CEA-binding protein 1 (CEABP1; Figures 3A and 3B; Figures S2A, S3A, and S3B), was predicted to possess high thermal stability, high binding affinity for CEA-A3, high solubility, and low immunogenicity, as verified by the epitope prediction tool in the Immune Epitope Database. The GFP-tagged CEABP1 protein was expressed in *E. coli*, purified, and was found via the immunofluorescence assay to bind to CEA on the cell membrane of LS174T CRC cells, which express high levels of CEA (Figure 3C). In addition, a binding protein for the CEA-B2 domain, CEABP2, was designed via RFDiffusion and ProteinMPNN (Figures 3D and 3E; Figures S2B, S3C, and S3D) and was confirmed to colocalize with CEA on the LS174T cell membrane (Figure 3F). The designed CEABP1 or CEABP2 was fused to the N- or C-terminal end of pep1-Max, and the resulting fusion proteins were expressed, purified, and used to deliver TCF/LEF TFD DNA into LS174T cells. In both the CCK-8 and colony formation assays, significantly greater suppression of CRC cell proliferation was obtained with the purified pep1-Max-CEABP1 protein than with pep1-Max alone. On the other hand, neither pep1-Max-CEABP2 nor CEABP2-pep1-Max convincingly showed a stronger ability to inhibit colorectal cancer cell growth than pep1-Max did (Figures 3G and 3H; Figure S4).

**Figure 2.**
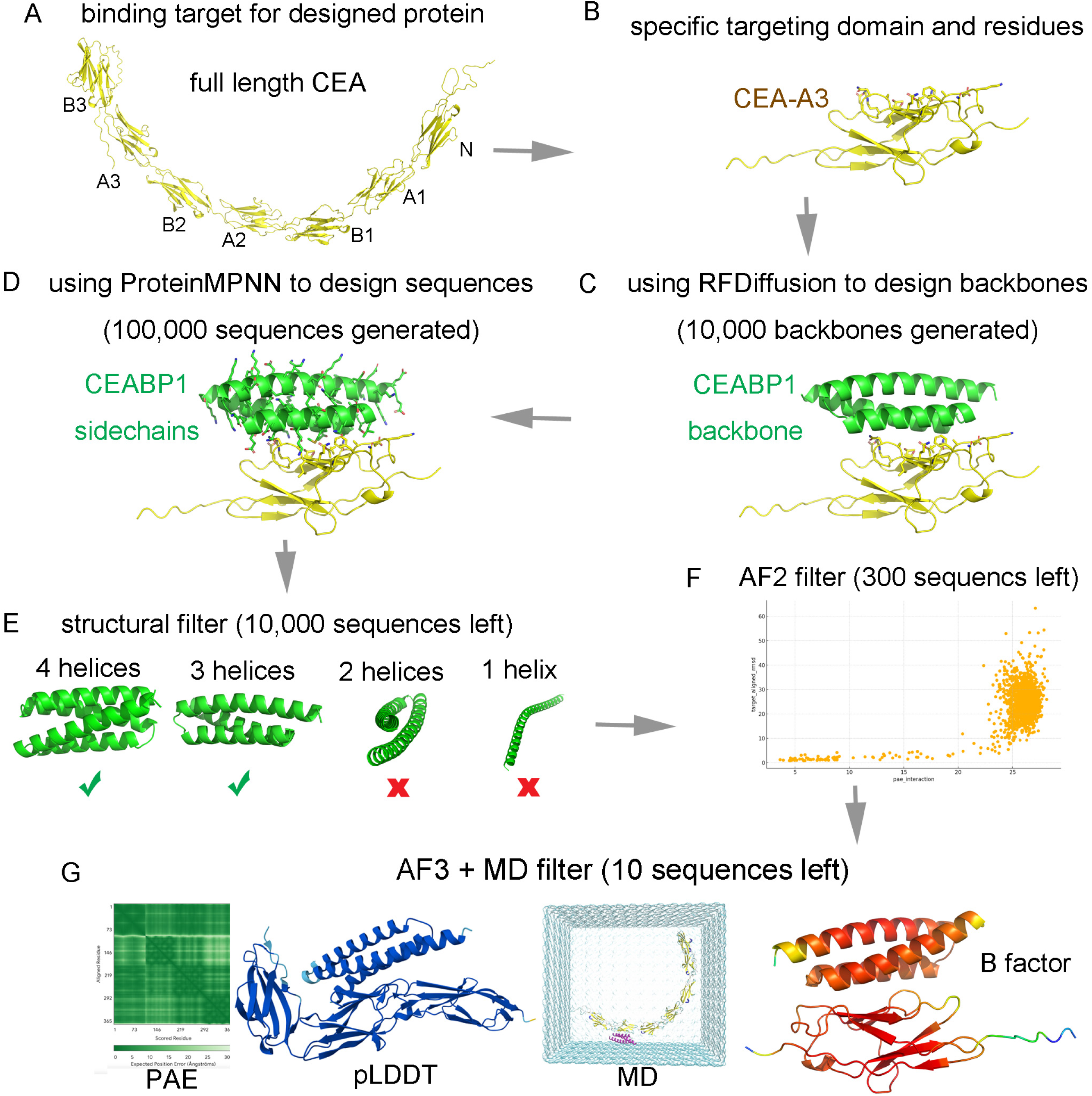
The strategy and workflow for the design of the binding protein CEABP1 for the A3 domain of CEA. (A) The structure of full length CEA was predicted by Alphafold3, which consisted of seven IgG domains. The available partial crystal structure of CEA is consistent with the predicted structure. (B) The A3 domain of CEA was selected for the specific targeting domain. The residues on one surface of CEA-A3 were chosen as “hotspots” for targeting binder design. (C) RFDiffusion was employed to design the backbone scaffolds of binding proteins for the CEA-A3 domain, during which process 10,000 backbones were generated. (D) ProteinMPNN was employed to design the sequences of these binding proteins, during which process 100,000 sequences were generated. (E) Generated binders were filtered based on the helix bundle topology. Designed proteins with four or three helices were selected, whereas those with two helices or only one helix were discarded. (F) Alphafold2 was employed to rapidly discard models that failed to recapitulate tight binding geometry with CEA-A3. (G) Molecular dynamics (MD) simulation for conformational sampling was performed, followed by Alphafold3 rescoring to identify false positives that passed Alphafold2 but were unstable during MD or had low predicted alignment error (PAE) or predicted local distance difference test (pLDDT) scores calculated by Alphafold3.

**Figure 3.**
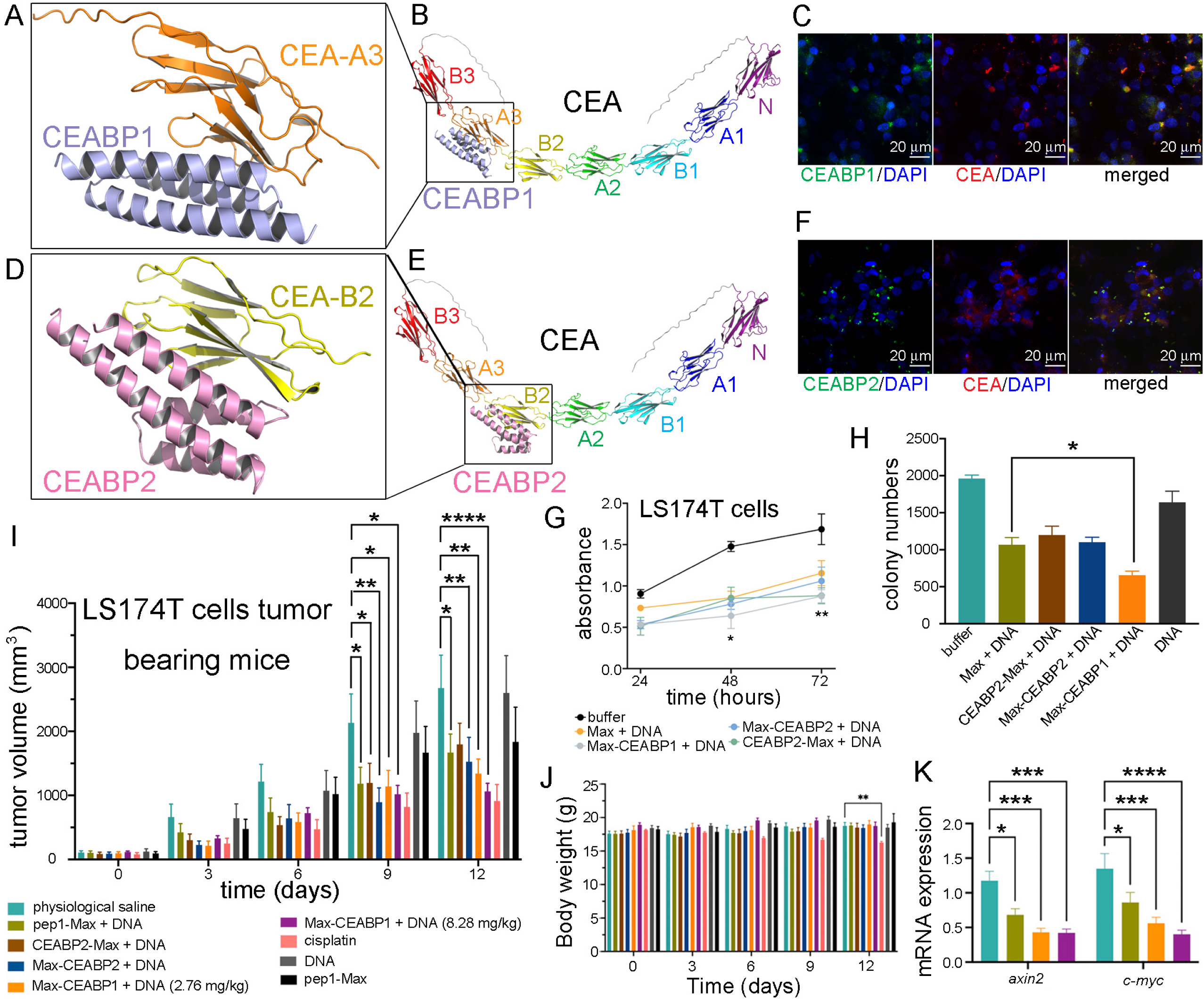
Specific targeting CEA-expressing CRC cells via a de novo-designed CEA-binding protein enhanced the inhibition of CRC cell proliferation and xenograft tumor growth by TCF/LEF TFD DNA. (A) Alphafold3-predicted structure of the extracellular A3 domain of CEA in complex with CEABP1, the de novo-designed binding protein for CEA-A3. (B) The binding site of CEABP1 on the full-length CEA protein. (C) CEABP1 colocalized with CEA on the cell membrane of the CEA-expressing CRC cell line LS174T. LS174T cells were incubated with purified GFP-tagged CEABP1 protein for 6 hours before immunofluorescence staining. Scale bars, 20 μm. (D) Alphafold3-predicted structure of the extracellular B2 domain of CEA in complex with CEABP2, the de novo-designed binding protein for CEA-B2. (E) The binding site of CEABP2 on the full-length CEA protein. (F) CEABP2 colocalized with CEA on the cell membrane of LS174T cells. (G) Fusion of CEABP1 to the C-terminus of pep1-Max enhanced the suppression of LS174T cell proliferation by the delivered TCF/LEF TFD DNA, which was examined via a CCK-8 assay. The data in the figure were compared between the pep1-Max-CEABP1 plus TCF/LEF TFD DNA group and the pep1-Max plus the same DNA group. (H) Colony formation assays also verified that fusing CEABP1 to pep1-Max caused the delivery of TCF/LEF TFD DNA to inhibit LS174T cell growth more strongly than the absence of CEABP1. (I) Targeting LS174T cell subcutaneous xenograft tumors with CEABP1 enhanced the ability of TCF/LEF TFD DNA to decelerate tumor growth in mice. The tumor growth curves of the following groups receiving different treatments are shown: physiological saline solution (n = 6), 2.76 mg/kg pep1-Max and TCF/LEF TFD DNA (n = 6), 2.76 mg/kg CEABP2-pep1-Max and TCF/LEF TFD DNA (n = 6), 2.76 mg/kg pep1-Max-CEABP2 and TCF/LEF TFD DNA (n = 6), 2.76 mg/kg pep1-Max-CEABP1 and TCF/LEF TFD DNA (n = 6), 8.28 mg/kg pep1-Max-CEABP2 and TCF/LEF TFD DNA (n = 6), 5 mg/kg cisplatin (n = 6), 7.34 mg/kg pep1-Max protein (n = 6), and 0.94 mg/kg TCF/LEF TFD DNA (n = 5). For the above protein and DNA mixtures, the protein-to-DNA molar ratio was 3:1. The error bars represent the SEMs, which were determined via two-way ANOVA. (J) Fusing CEABP1 or CEABP2 to pep1-Max did not cause any body weight changes in the mice during the experiment. (K) Specific targeting CEA-expressing LS174T cells with CEABP1 further decreased the mRNA expression of Wnt-responsive genes in tumors. qRT□PCR analysis of *axin2* and *c-myc* mRNA transcript levels in tumor samples from the indicated groups was performed.

To investigate whether the use of CEABP1 or CEABP2 to specifically target CEA-expressing CRC cells could increase the tumor growth inhibition efficacy of TCF/LEF TFD DNA, an in vivo study in immunocompromised athymic nude mice bearing LS174T xenografts was performed. TCF/LEF TFD DNA, together with purified pep1-Max, CEABP2-pep1-Max, pep1-Max-CEABP2, or pep1-Max-CEABP1 proteins, were systemically injected via the tail vein every three days for four successive treatments. LS174T tumor-bearing mice treated with physiological saline, pep1-Max protein alone, or TCF/LEF TFD DNA alone displayed rapid tumor growth, whereas those treated with TCF/LEF TFD DNA in combination with pep1-Max-CEABP1 presented more decelerated tumor growth than those treated with pep1-Max, CEABP2-pep1-Max, or pep1-Max-CEABP2 (Figure 3I; Figure S5). Nevertheless, there was no perceivable effect on mouse body weight when mice were treated with a TCF/LEF TFD together with CEABP1- or CEABP2-fused pep1-Max (Figure 3J). To evaluate whether specifically targeting CRC cells indeed strongly inhibited Wnt signaling, mRNA from LS174T tumors in different treatment groups was extracted and analyzed via qRT□PCR. As shown in Figure 3K, the use of pep1-Max-CEABP1 to target CEA-expressing CRC cells and deliver TCF/LEF TFD DNA indeed enhanced the suppression of Wnt responsive genes such as *axin2* and *c-myc* in the tumor tissue in comparison with the use of pep1-Max.

Therefore, the use of a fusion protein of the de novo-designed CEA-binding protein CEABP1 and pep1-Max to specifically target CRC cells further enhanced the suppression of CRC cell proliferation and xenograft tumor growth by TCF/LEF TFD DNA compared with the use of pep1-Max for delivery.

### Delivery of the purified p53 protein with the cell-penetrating peptide p28 suppressed colorectal cancer cell proliferation and xenograft tumor growth

In CRC, mutations not only occur in the Wnt signaling pathway but also in p53, which occurs in 60% of nonhypermutated CRC cases. p53 functions as a master regulator of cell growth and proliferation by mediating cell cycle arrest, apoptosis, senescence, metabolism, ferroptosis, and immunity. The protein level of p53 is usually kept low by E3 ubiquitin ligases, including Mdm2, ARF-BP1/Mule, and COP1.^14^

We explored the feasibility of using cell-penetrating peptides to deliver purified p53 protein into CRC cells. p53 proteins fused with different CPPs, including pep1, penetratin (pene), p28, or TAT, were expressed and purified to homogeneity (Figure 4A) and then examined via HCT116 cell colony formation assay (Figure 4B; Figure S6). Among the different CPP-p53 proteins, p28-p53 exhibited the strongest suppression of CRC cell proliferation and xenograft tumor growth. In addition, in the HCT116 xenograft mouse tumor growth assay, purified p28-p53 protein, but not purified p53 protein without p28, exhibited potent inhibition of xenograft tumor growth (Figure 4C).

**Figure 4.**
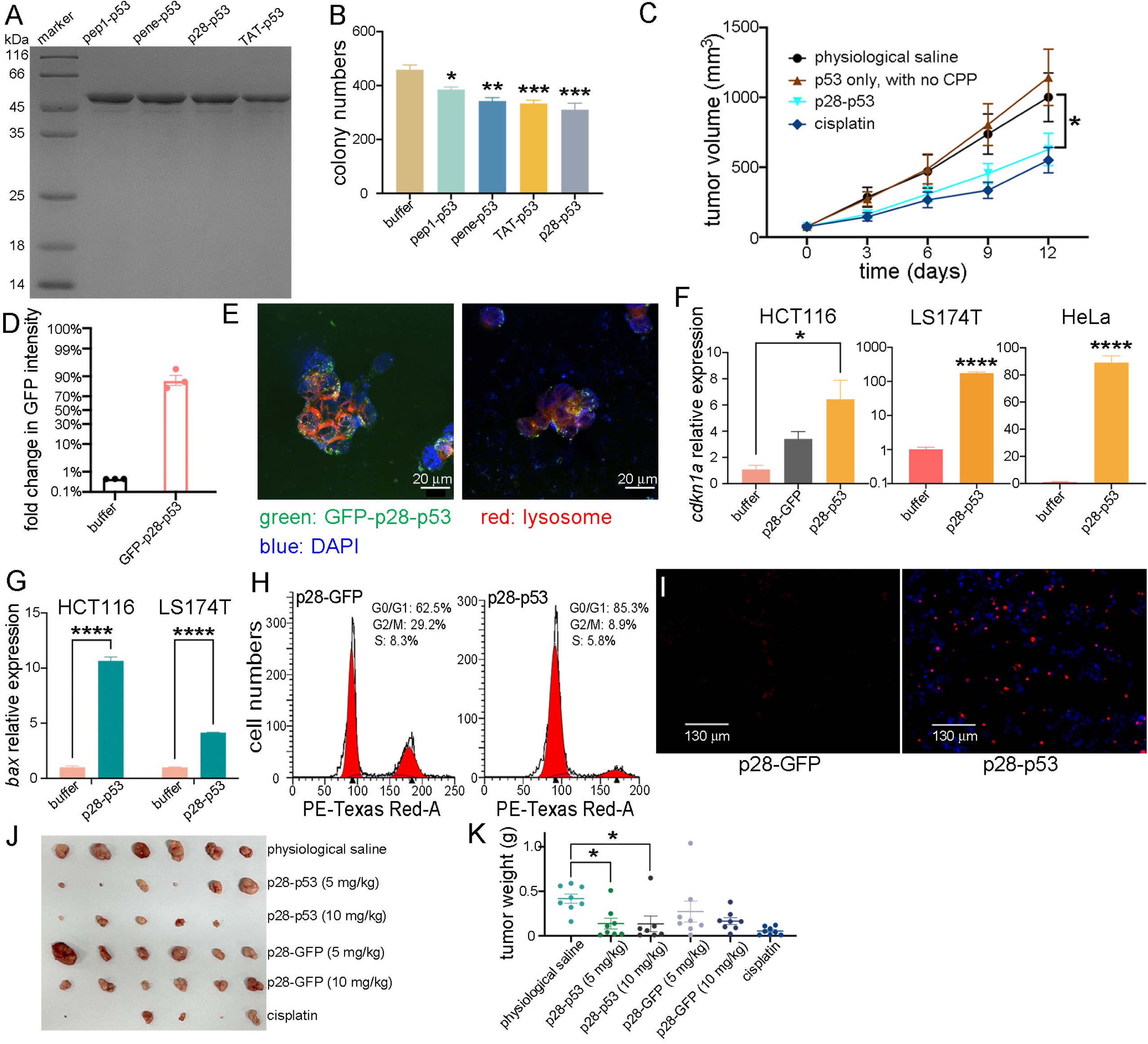
Delivery of purified p53 protein via the cell-penetrating peptide p28 suppressed CRC cell proliferation and xenograft tumor growth in mice. (A) Purification of the full-length human p53 protein fused with various cell-penetrating peptides: pep1, penetration (pene), p28, and TAT. (B) Purified p28-p53 protein exhibited the strongest inhibition of cell proliferation when delivered into HCT116 cells, as shown by the colony formation assay. (C) Purified p28-p53 protein most effectively restrained the growth of HCT116 cell subcutaneous xenograft tumors in mice. The tumor growth curves of the following groups receiving different protein delivery treatments are shown: physiological saline solution (n = 7), 10 mg/kg p53 without CPP (n = 8), 10 mg/kg p28-p53 (n = 8), and 5 mg/kg cisplatin (n = 8). (D) Purified GFP-tagged p53 protein could be highly efficiently delivered into HCT116 cells via the p28 CPP, as suggested by the fold change in GFP fluorescence intensity after protein delivery. (E) Confocal microscopy images showing that after delivery into HCT116 cells, the internalized p28-GFP-p53 protein escaped from lysosomes and was localized in the nucleus. (F) Delivery of the p28-p53 protein effectively increased *cdkn1a* transcript levels in HCT116, LS174T, and HeLa cells, as revealed by qRT□PCR analysis. (G) Delivery of the p28-p53 protein also increased the transcript level of *bax* in HCT116 and LS174T cells, as shown by qRT□PCR analysis. (H) Delivery of the p28-p53 protein caused cell cycle arrest at the G0/G1 phase in HCT116 cells. The cell cycle distribution was determined via flow cytometry. (I) After delivery, p28-p53 protein triggered apoptosis in HCT116 cells, as revealed by the TUNEL assay. The red dots represent apoptotic cells. (J) Treatment of HCT116 cell subcutaneous xenograft tumor-bearing mice with purified p28-p53 protein decreased tumor growth. Representative images of HCT116 cell xenograft tumors at the endpoint of the experiment are shown: physiological saline solution (n = 6), 5 mg/kg p28-p53 (n = 6), 10 mg/kg p28-p53 (n = 5), 5 mg/kg p28-GFP (n = 6), 10 mg/kg p28-GFP (n = 6), and 5 mg/kg cisplatin (n = 5). (K) The weight of HCT116 cell xenograft tumors at the endpoint.

p28 is a CPP derived from *Pseudomonas aeruginosa* that can preferentially enter cancer cells through endocytosis and interfere with the E3 ubiquitin ligase COP1.^39,40,41^ We further confirmed that the purified GFP-tagged p28-p53 protein could be efficiently delivered into HCT116 CRC cells, with a fold change in the GFP intensity of approximately 90% (Figure 4D). In addition, the delivered GFP-p28-p53 protein escaped from lysosomes and localized to the nuclei of HCT116 cells (Figure 4E). qRT□PCR analysis demonstrated that the delivered p28-p53 protein indeed increased the expression of p53 target genes such as *cdkn1a* (Figure 4F) and *bax* (Figure 4G) in both HCT116 and LS174T CRC cells. Additionally, after delivery of the purified p28-p53 protein, G0/G1 phase cell cycle arrest was triggered (Figure 4H), and apoptosis was promoted (Figure 4I), presumably as a result of elevated p21^Cip^^1^ and Bax protein levels. To further validate the therapeutic efficacy of the purified p28-p53 protein, an in vivo study using immunocompromised athymic nude mice bearing HCT116 cell xenografts was performed. Intravenous injection of purified p28-p53 protein at two different doses, 5 mg/kg and 10 mg/kg, every 3 days for the six treatments both decelerated CRC cell xenograft tumor growth (Figure 4J) and substantially decreased tumor weight at the experimental endpoint (Figure 4K).

In summary, these data demonstrated that the delivery of purified p28-p53 protein effectively inhibited CRC cell proliferation and suppressed xenograft tumor growth in mice.

### Delivery of purified p28-p14^ARF^ (residues 1-63) protein prolonged the effective duration of delivered p53 protein and increased its antitumor effect

p14^ARF^ is another crucial tumor suppressor protein that prevents Mdm2 from downregulating p53 by regulating its nucleocytoplasmic shuttling and ubiquitination.^20–24^ In addition, it has been reported that p14^ARF^ directly binds and inhibits another E3 ubiquitin ligase, ARF-BP1/Mule.^19^ Thus, the possibility of using delivered p14^ARF^ protein to prolong the effective duration of delivered p53 protein in CRC cells was explored. The crystal structure of p14^ARF^ in complex with Mdm2 has not yet been reported, despite the continuous efforts of multiple laboratories. Therefore, Alphafold3 was used to obtain a predicted human p14^ARF^-Mdm2 complex structure, which showed that the N-terminal 63 residues of p14^ARF^ folded into two β-strands and an α-helix and interacted with a middle fragment of Mdm2, residues 186-255 (Figure 5A). The predicted structure of the complex between p14^ARF^ (1-63) and Mdm2 (186-255) was shown to remain stable during the molecular dynamics simulation process. The β1 strand of p14^ARF^ formed main-chain hydrogen bond interactions with a β-strand from Mdm2 (residues 246-254). In addition, hydrophobic interactions dominated at the center of the p14^ARF^-Mdm2 interface, and electrostatic interactions between the positively charged arginines of p14^ARF^ and the negatively charged residues of Mdm2 existed at the periphery of the binding interface (Figure S7). Importantly, residues 1-63 of p14^ARF^ exhibited extensive conservation among representative species from various orders within mammals (p14^ARF^ only exists in mammals), suggesting that these residues may also be critical for other functions of p14^ARF^, such as inhibiting ARF-BP1/Mule (Figure S8). Therefore, the N-terminal domain of p14^ARF^ (residues 1-63, abbreviated as p14^ARF^ hereafter) was employed in the following study.

**Figure 5.**
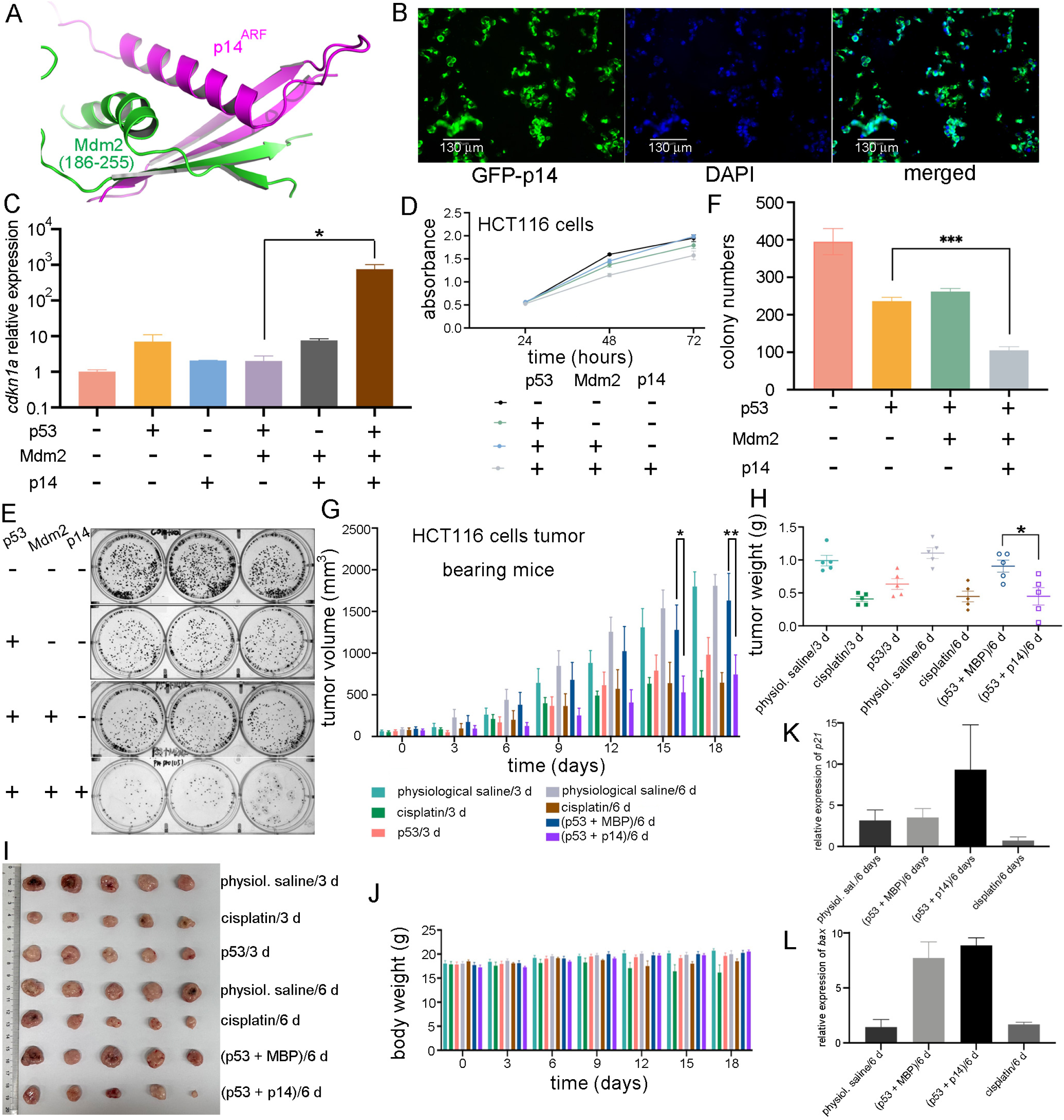
Delivery of the purified p28-p14^ARF^ protein prolonged the effective duration of the delivered p53 protein and strengthened its anti-proliferative and antitumor effects. (A) The N-terminal 63 residues of p14^ARF^ were predicted by Alphafold3 to fold into two β strands and an α helix and interact with residues 186-255 of human Mdm2. (B) Purified GFP-tagged p28-p14^ARF^ (residues 1-63, abbreviated as p28-p14^ARF^) protein was localized in the nucleus after delivery into HCT116 cells. Scale bars: 130 μm. (C) Delivery of the p14^ARF^ protein counteracted the inhibition of p53 by Mdm2 and drastically increased the expression of the p53 target gene *cdkn1a*. Purified p28-p53 and p28-p14^ARF^ proteins were delivered while an Mdm2-encoding plasmid was transfected into HCT116 cells, and qRT□PCR analysis was performed. (D) The delivered p14^ARF^ protein enhanced the inhibitory effect of p53 on CRC cell proliferation, as shown by the CCK-8 assay. Purified p28-p14^ARF^ and p28-p53 proteins were delivered while an Mdm2-encoding plasmid was transfected into HCT116 cells. (E) Colony formation assays revealed that the purified p14^ARF^ protein further increased the anti-proliferative effect of the purified p28-p53 protein on HCT116 cells. (F) Quantification of the colony formation assay results from (E). (G) Compared with the injection of p28-p53 and MBP proteins once every 6 days, the injection of p28-p53 and p28-p14^ARF^ proteins once every 6 days substantially reduced the tumor growth rate in HCT116 cell subcutaneous xenograft mice. The tumor growth curves of the following groups receiving different treatments are shown: physiological saline solution every 3 days (n = 5), 5 mg/kg p28-p53 every 3 days (n = 5), 5 mg/kg cisplatin every 3 days (n = 5), physiological saline solution every 6 days (n = 5), 5 mg/kg p28-p53 and p28-MBP each every 6 days (n = 5), 5 mg/kg p28-p53 and p28-p14^ARF^ each every 6 days (n = 5), and 5 mg/kg cisplatin every 6 days (n = 7). (H) The corresponding HCT116 cell xenograft tumor weight at the endpoint. (I) Representative images of HCT116 cell xenograft tumors at the endpoint of the experiment (day 18). (J) Treatment with purified p28-p53 and p28-p14^ARF^ proteins did not cause body weight changes in the mice during the experiment. The color scheme for the different groups was the same as that described in (G). (K) qRT□PCR analysis revealed that treatment with purified p28-p53 and p28-p14^ARF^ proteins every 6 days increased the mRNA expression of *cdkn1a* in the tumor, whereas treatment with purified p28□p53 and MBP proteins every 6 days had no effect. (L) qRT□PCR analysis revealed that treatment with purified p28-p53 and p28-p14^ARF^ proteins every 6 days increased the mRNA expression of *bax* in tumors.

The purified GFP-tagged p28-p14^ARF^ protein was found to be localized in the nucleus after delivery into HCT116 cells (Figure 5B). To verify that the delivered p28-p14^ARF^ protein could antagonize the inhibitory effect of Mdm2 on p53, a qRT□PCR assay was performed, which revealed a dramatic increase in *cdkn1a* expression due to the delivery of p53 when the p28-p14^ARF^ protein was simultaneously delivered (Figure 5C). Furthermore, both the CCK-8 (Figure 5D) and colony formation assay (Figures 5E and 5F) results suggested that delivery of purified p28-p14^ARF^ protein enhanced the anti-proliferative effect of the delivered p53 protein in HCT116 cells, presumably as a result of inhibition of Mdm2 and ARF-BP1/Mule and subsequent stabilization of p53.

Immunocompromised athymic nude mice bearing HCT116 xenografts were injected every 6 days with purified p28-p53 and p28-p14^ARF^ proteins or with purified p28-p53 and maltose-binding protein (MBP) proteins as a control. Administration of the p28-p53 and MBP proteins every 6 days did not slow down the progression of HCT116 xenograft tumors, in contrast to treatment with the p28-p53 protein every 3 days, which is likely due to the short half-life of the p53 protein in cells resulting from the negative regulation of E3 ubiquitin ligases. On the other hand, with codelivery of the p28-p14^ARF^ protein, extending the dosing interval of p28-p53 to six days resulted in considerable inhibition of subcutaneous HCT116 tumor growth in the mice (Figures 5G and 5H). There was a statistically significant difference between the tumor weights of the mice treated with p53 and p14^ARF^ and those of the mice receiving p53 and MBP (Figure 5I). Moreover, treatment with the p28-p14^ARF^ and p28-p53 proteins did not affect the body weights of the mice (Figure 5J). Consistent with the findings in HCT116 cell culture, p28-p14^ARF^ protein codelivery significantly enhanced p53-mediated upregulation of the downstream target genes *cdkn1a* (Figure 5K) and *bax* (Figure 5L) in tumor tissue.

In conclusion, codelivery of the purified p28-p14^ARF^ protein with p53 substantially enhanced the antiproliferative effect of the delivered p53 protein in CRC cells, and simultaneous treatment of mice with the purified p28-p14^ARF^ protein enhanced the antitumor effect of the delivered p53 protein by prolonging its effective duration.

### Specifically targeting CRC cells with p28-p53-CEABP1 resulted in considerable higher suppression of CRC cell proliferation and xenograft tumor growth

In light of the success of using a fusion protein of a de novo-designed CEA-binding protein and pep1-Max to specifically target CRC cells for delivery of TCF/LEF TFD DNA and suppression of CRC cell proliferation, the CEABP1 or CEABP2 protein was fused either to the N- or C-terminus of the p28-p53 protein. With the exception of the C-terminal fusion of CEABP2 to p28-p53, which failed to yield detectable protein expression, the CEABP1-p28-p53, p28-p53-CEABP1, and CEABP2-p28-p53 proteins were purified to homogeneity and delivered to the CEA-expressing LS174T CRC cell line to examine the transcription of p53 target genes via qRT□PCR. Among the three fusion proteins examined, the delivery of the purified p28-p53-CEABP1 protein significantly increased the transcription of the p53 target genes *cdkn1a* (Figure 6A) and *bax* (Figure 6B) in LS174T cells compared with the delivery of p28-p53. In addition, delivery of the p28-p53-CEABP1 protein resulted in even stronger suppression of LS174T cell proliferation than delivery of the p28-p53 protein in the CCK-8 assay (Figure 6C). To further validate the tumor-suppressive effect of p28-p53-CEABP1 in vivo, we established a subcutaneous LS174T xenograft tumor model in nude mice. Consistent with the cell assay results, tail vein injection of the p28-p53-CEABP1 protein demonstrated substantially greater inhibitory efficacy (p < 0.0001) against LS174T tumor growth than did p28-p53 protein treatment (Figures 6D and 6E). At the end of the experiment, the tumor weight of the group of mice receiving p28-p53-CEABP1 protein was considerably lower than that of the group receiving p28-p53 treatment (Figure 6F), whereas the body weights of the mice did not seem to be affected throughout the experiment (Figure 6G). In addition, hematoxylin and eosin staining analysis of harvested major organs (heart, lung, liver, kidney, and spleen) revealed no detectable histopathological abnormalities in the mice that received p28-p53, p28-p53 together with p14, or p28-p53-CEABP1 treatment (Figure S9). The inhibitory effect on CRC cell xenograft tumor growth was demonstrated to be mediated through elevated p53-regulated expression of the cell cycle inhibitor p21^Cip^^1^ and the proapoptotic protein Bax in tumors, as evidenced by qRT□PCR analysis of tumor tissue samples (Figures 6H and 6I). TUNEL assay results using tumor tissue samples also revealed that apoptosis was further elevated by treatment with the delivered p28-p53-CEABP1 protein than by treatment with p28-p53 (Figure 6J).

**Figure 6.**
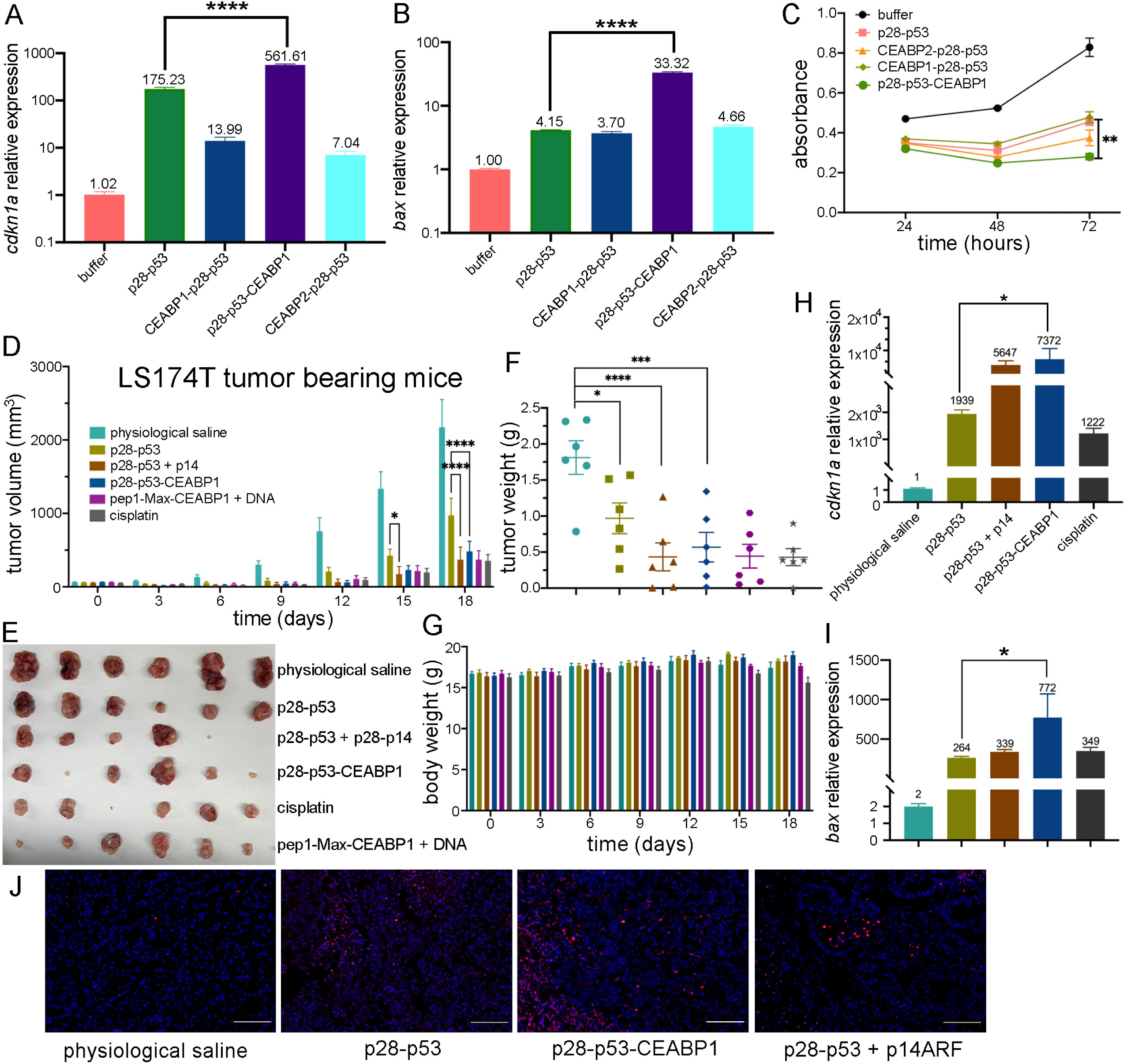
Specific targeting CRC cells by fusing CEABP1 to the C-terminus of p28-p53 considerably enhanced its ability to suppress CRC cell proliferation and xenograft tumor growth in mice. (A) qRT□PCR analysis revealed that fusing CEABP1 to the C-terminus of p28□p53 was most effective at increasing the ability of the delivered p53 protein to promote *cdkn1a* transcription in LS174T cells. The relative *cdkn1a* expression levels are indicated above the columns. (B) The p28-p53-CEABP1 protein was most effective at increasing the transcription of *bax* when it was delivered into LS174T cells. The relative *bax* expression levels are indicated above the columns. (C) Compared with delivery of p28-p53, delivery of the purified p28-p53-CEABP1 protein more strongly inhibited the proliferation of LS174T cells, as shown by the CCK-8 assay. (D) Treatment with the purified p28-p53-CEABP1 protein inhibited LS174T cell subcutaneous xenograft tumor growth more effectively than did p28-p53. The tumor growth curves of the following groups receiving different treatments are shown: physiological saline solution (n = 6), 5 mg/kg p28-p53 (n = 6), 5 mg/kg p28-p53 and 5 mg/kg p28-p14^ARF^ (n = 6), 5 mg/kg p28-p53-CEABP1 (n = 6), 8.28 mg/kg p28-p53-CEABP1 and TCF/LEF TFD DNA (molar ratio 3:1, n = 6), and 5 mg/kg cisplatin (n = 6). (E) Representative images of LS174T cell xenograft tumors at the endpoint (day 18). (F) LS174T tumor weight at the endpoint. The color scheme for the different groups was the same as that in (D). (G) Treatment with p28-p53-CEABP1 did not cause perceptible changes in mouse body weight during the experiment. The color scheme for the different groups was the same as that described in (D). (H) qRT□PCR analysis revealed that, compared with the absence of CEABP1, the injection of purified p28-p53-CEABP1 substantially increased the *cdkn1a* transcript level in the tumor tissue. (I) qRT□PCR analysis revealed that p28-p53-CEABP1 drastically increased *bax* mRNA expression compared with p28-p53 in tumors. The color scheme for the different groups is the same as that in (H). (J) Fluorescence microscopy images of fixed tumor tissues subjected to TUNEL staining. Blue: DAPI; red: apoptotic cells. Scale bars: 100 μm.

In summary, fusing the CEABP1 protein to the C-terminus of p28-p53 for specific targeted delivery to CRC cells substantially enhanced the ability of the delivered p53 protein to inhibit CRC cell proliferation and xenograft tumor growth.

## Discussion

While previous efforts in the development of anticancer therapeutics based on interference with Wnt signaling have focused on the critical step of the association between β-catenin and TCF/LEF, our work endeavored to target the subsequent step of recognition between TCF/LEF and its target gene promoters. Our study demonstrated that TCF/LEF transcription factor decoy (TFD) DNA conjugated with the pep1-Max-CEABP1 protein could effectively suppress Wnt signaling in CRC cells ex vivo and in vivo, potentially offering a promising strategy to regulate Wnt-driven processes with enhanced specificity and reduced off-target toxicity. Our therapeutic strategy directly targeted the most downstream step of the Wnt signaling pathway and could counteract any kind of activating mutation in this pathway. The choice of the pep1-Max-CEABP1 protein as a DNA delivery mediator was crucial, as it not only provided efficient nuclear delivery of TFD DNA but also ensured CRC cell-targeting specificity. This approach circumvents the limitations of traditional TFD DNA delivery methods, which suffer from low cellular uptake, improper subcellular localization, and poor targeting specificity for cancer cells. Importantly, the significant reduction in tumor volume in the xenograft model highlighted the potential of this strategy to directly modulate oncogenic pathways in CRC. Several noteworthy issues in the choice of DNA are as follows. (i) The consensus binding motif of TCF/LEF, 5□-AGATCAAAGG-3□, is relatively long. Therefore, high targeting specificity for TCF/LEF was ensured, and the possibility of off-target binding to other transcription factors was kept low. (ii) The immune response triggered by the DNA-activated cGAS/STING pathway is critically related to the length of the DNA, and that induced by DNA shorter than 45 bp has been shown to be weak.^46,47^ In this work, 36 bp TFD DNA was used. (iii) The ends of the TFD DNA used were modified by phosphorothioation to prevent exonucleases from degrading the delivered DNA.

Therapeutic strategies targeting p53 present both unprecedented opportunities and formidable challenges in oncology. Although small-molecule Mdm2 inhibitors such as nutlin have been developed to inhibit p53 ubiquitination by Mdm2, these Mdm2 inhibitors may be effective only for patients with wild-type *TP53* genes and may not work for cancer patients with mutated *TP53*. Ways have also been developed to restore the activity of mutated p53 protein, using arsenic trioxide (ATO) to bind to allosteric sites on p53 undergoing structural mutations, for example.^48^ However, this approach was not effective for large numbers of DNA-contacting mutants of p53, such as R273H.^49^ While mutant p53 reactivators such as APR-246 partially restored p53 transcriptional activity in *TP53*-mutant myelodysplastic syndromes, the overall response rates remained modest, with significant interpatient heterogeneity.^50^ While current precision oncology frameworks emphasize mutation-specific reactivation of p53, clinical realities reveal staggering heterogeneity resulting from *TP53* mutation spectra. In this work, CRC cell-specific restoration of purified recombinant p53 protein was accomplished via the use of p28 for delivery, a CPP that also inhibits the E3 ubiquitin ligase COP1, and an artificially designed CEABP1 protein module for targeting. Furthermore, codelivery of the purified p28-p14^ARF^ protein significantly prolonged the effective duration of the p53 protein by inhibiting MDM2- and ARF-BP1/Mule-mediated ubiquitination and degradation. This approach not only enhances antitumor effects but also reduces the dosing frequency in mouse models. Equally noteworthy is that this method could engage erroneous p53 signaling pathways regardless of mutation type.

The integration of cutting-edge computational tools for de novo protein design was pivotal for ensuring CRC-specific targeting in our work. By recognizing CEA, a well-characterized CRC biomarker localized on the cell membrane, the designed CEABP1 protein enhanced the selectivity of TCF/LEF TFD and p53 protein delivery to CEA-expressing tumor cells. The increased antitumor activity observed with pep1-Max-CEABP1 or p28-p53-CEABP1 over nontargeted controls pep1-Max or p28-p53, respectively, underscores the effectiveness of targeted delivery. This finding aligns with the success of antibody□drug conjugates (ADCs) in clinical oncology, suggesting that protein-based drugs could serve as versatile platforms for precision cancer therapy. Such workflows can be adapted to target other cancer-associated biomarkers, expanding the applicability of protein-based therapeutics.

While the present results are encouraging, several challenges remain. The long-term effectiveness and biodistribution of the delivered DNA and p53 protein in vivo require optimization. Additionally, the potential immunogenicity of synthetic proteins must be reduced to a minimum. There are twenty lysines in the wild-type p53 protein, which are potential ubiquitination sites for at least five kinds of E3 ubiquitin ligases. An engineered or redesigned p53-like protein in which most or all lysines are replaced would be exempt from ubiquitination and degradation and would have a further elongated half-life. In addition, the thermal stability of the DNA-binding domain of p53 can hardly be considered superior, and a reengineered DNA-binding domain would also benefit from tighter folding and enhanced stability. In addition, TCF/LEF TFD DNA or p53-based therapy could be combined with existing CRC treatments (e.g., chemotherapy or immunotherapy) to obtain synergistic effects, and the combination of the modulation of Wnt signaling and p53 restoration might also enhance therapeutic efficacy. Our studies could also be extended to investigate the targeting of other oncogenic transcription factors by TFD DNA and the delivery of other tumor suppressor proteins, such as Rb or p16^INK4a^, via elaborately designed tumor marker recognition proteins for precision cancer therapeutics.

## Methods

### Protein expression and purification

All the plasmids were constructed via homologous recombination. Genes encoding various cell-penetrating peptides, including p28, TAT, penetratin (pene), pep1, and the TCF transcription factor decoy DNA, and genes encoding the designed CEA-binding proteins (including CEABP1 and CEABP2) were synthesized by the BGI company. The pMAL-C2x vector was used to express the p28-p53, TAT-p53, pene-p53, pep1-p53, p28-p53-CEABP1, and p28-CEABP2-p53 proteins. The modified pET28a vector with the MBP tag added was used to express the pep1-Max, pep1-Max-CEABP1, pep1-Max-CEABP2, and CEABP2-pep1-Max proteins. The pET28a vector was used to express the GFP-CEABP1 and GFP-CEABP2 proteins. The plasmids were subsequently transformed into *E. coli* BL21(DE3) cells. For protein overexpression, 10 mL of overnight culture was inoculated into 1 L of LB medium supplemented with 100 mg/mL ampicillin or kanamycin. The culture was induced with 0.2 mM IPTG and further incubated at 16°C for 16 hours. The cells were then centrifuged, resuspended in MBP affinity column binding buffer (25 mM Tris–HCl, pH 8.0, 300 mM NaCl, and 2 mM DTT) or Ni^2+^-NTA column binding buffer (25 mM Tris–HCl, pH 8.0, 300 mM NaCl, and 20 mM imidazole), and lysed by a cell homogenizer (JNBio) at 4°C. After centrifugation (18,000 rpm for 30 min at 4°C), the supernatant was applied to an MBP affinity column (Smart-Life Sciences, China) and eluted with 20 mM maltose or applied to a Ni^2+^-NTA affinity column (QIGEN) and eluted with a 40–500 mM linear concentration gradient of imidazole. The MBP-tagged proteins were cleaved by TEV protease at 4°C overnight. Finally, the proteins above were further purified by gel filtration chromatography using a Superdex 200 GL 10/300 column (GE Healthcare), using phosphate-buffered saline (PBS), pH 7.4, or physiological saline solution. The peak fractions were combined and concentrated to 2 mg/mL. The purified proteins were analyzed via 12% SDS–PAGE and Coomassie blue staining. Protein concentrations were determined via a Bradford protein assay kit (Bio-Rad).

### Cell lines and cell culture

The cells used in this study included (i) the human colorectal cancer cell line HCT116 (ATCC#CCL-247^TM^), with wild-type p53; (ii) the human colorectal cancer cell line SW480 (ATCC#CCL-228^TM^), with two point mutations of p53 (R273H/P309S); (iii) the human colorectal cancer cell line LS174T (ATCC#CCL-188^TM^), with wild-type p53; and (iv) the human cervical cancer cell line HeLa (ATCC#CCL-2^TM^). All cells were grown in Dulbecco’s modified Eagle’s medium (DMEM, Corning) supplemented with 100 U/ml penicillin, 100 mg/ml streptomycin (Gibco), 20 mM L-glutamine and 10% fetal bovine serum (FBS; Bovogen Biologicals, Australia).

### Quantitative real-time reverse transcription polymerase chain reaction (qRT□PCR)

qRT□PCR was used to quantify the expression of the p53 target genes *cdkn1a* (encoding the p21^Cip^^1^ protein) and *bax*, as well as the Wnt signal transduction pathway target genes *cyclin D1* and *c-myc* in the HCT116, SW480, LS174T, and HeLa cell lines. The cells were plated in 24-well plates at a density of 5×10^5^ cells per well. After 24 hours of cell adherence, protein or DNA delivery into the cells was performed via the addition of 20 μg of protein or 1 ng of DNA per well for 6 hours. After delivery, 1 mL of fresh complete medium was added, and the cells were further incubated for another 24 or 48 hours before RNA extraction and qRT□PCR. Total RNA was isolated via an RNA Extraction Kit (SPARKeasy) according to the manufacturer’s protocol. The RNA was quantified by measuring the UV absorbance at 260 nm. Next, the cDNA was reverse transcribed via a cDNA synthesis kit, PrimeScript™ RT Master Mix Perfect Real Time (Takara). qRT□PCR was performed in a real-time PCR detection instrument with SYBR Green dye (CWBIO). A total of 20 μL of mixture containing 100 ng of cDNA, 2 μM forward and reverse primers each, and 10 μL of 2×MagicSYBR mixture was used for each PCR. The fluorescence signal was recorded at the endpoint of each cycle during the 40 cycles (denaturing at 95°C for 15 seconds and annealing at 60°C for 30 seconds). *gapdh* was used as an internal control gene for normalization. Relative gene expression was calculated via the 2^−ΔΔCt^ method, which represents the inverse of the amount of mRNA in the initial sample.

The sequences of primers used for qRT□PCR are listed below. Data were analyzed by qPCRsoft

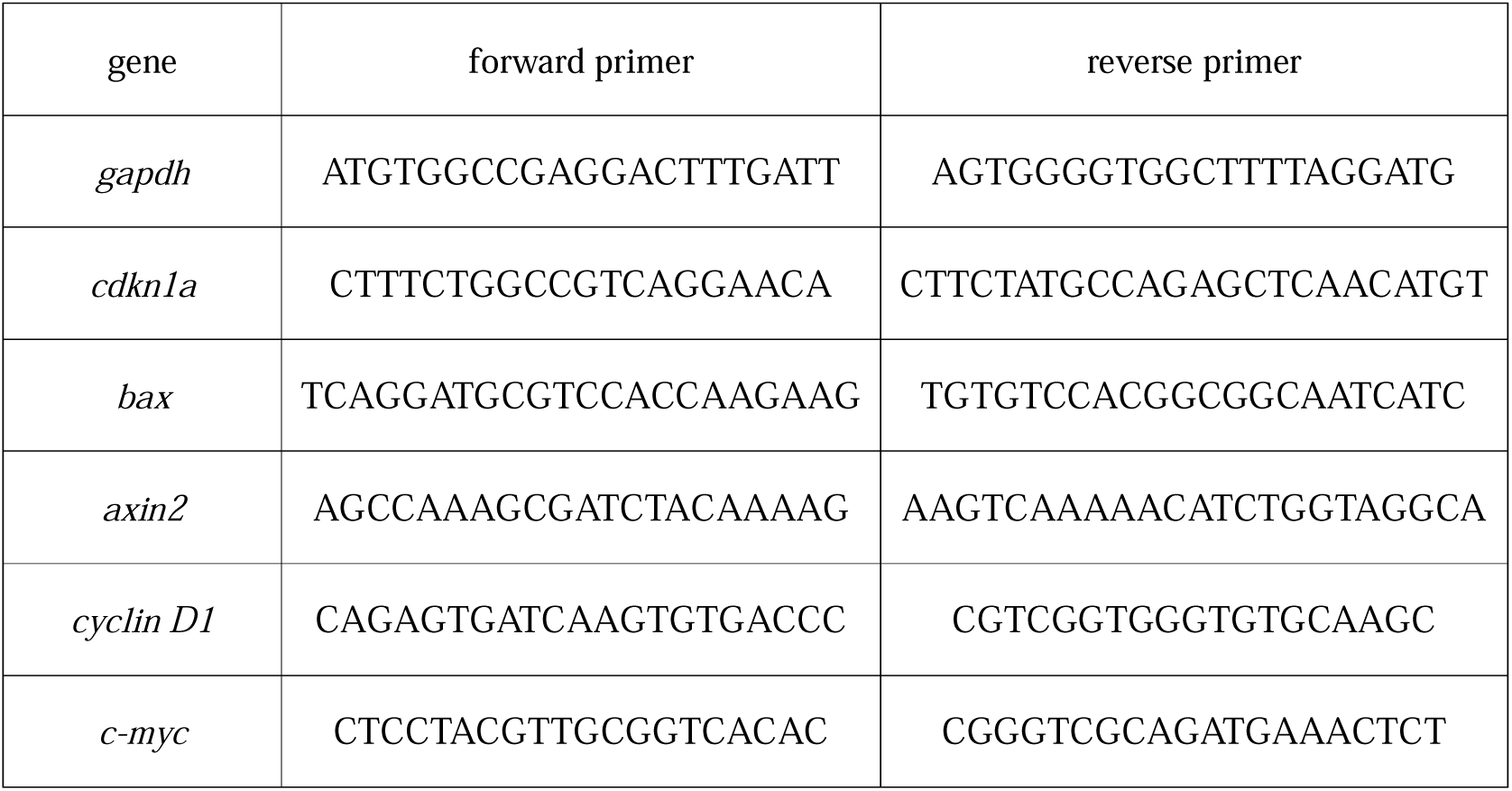

### CCK-8 cell proliferation assay

Colorectal cancer cell proliferation was examined via the Cell Counting Kit-8 (CCK-8) assay (Meilunbio, Dalian, China). The cancer cells were plated in 24-well plates at a density of 5×10^5^ cells per well. After 24 hours of cell adherence, protein or DNA delivery into cells was performed via the addition of 20 μg of protein or 1 ng of DNA per well for 6 hours. Afterwards, the cells were detached with 0.25% EDTA-treated trypsin and seeded into 96-well plates at a density of 3×10^3^ cells per well. After 24, 48, or 72 hours, the old medium was discarded, and 0.1 mL of new medium without FBS but containing 0.01 mL of CCK8 solution was added to each well and incubated for 0.5–4 hours. The absorbance was determined at a wavelength of 450 nm via a microplate reader.

### Colony formation assay

The cells were treated with purified p53 protein or DNA for 48 hours. Then, the cells were detached with 0.25% EDTA-treated trypsin, seeded into 6-well plates at a low density (∼1000 cells per well), and incubated for 2 weeks. The plates were then washed with PBS, fixed in 4% paraformaldehyde for 20 minutes, and then stained with 0.005% crystal violet. The images of all the wells were scanned and analyzed.

### Terminal deoxynucleotidyl transferase-mediated dUTP-biotin nick end labeling (TUNEL) assay

Apoptotic cells were measured with a One Step TUNEL Apoptosis Assay Kit (Beyotime) according to the manufacturer’s protocol. Tumors were extracted and fixed in formalin, embedded in paraffin, and sectioned at a thickness of 5 μm. TUNEL-positive cells had pyknotic nuclei with red fluorescence, indicating apoptosis.

Images of the sections were taken with a fluorescence microscope (Echo).

### Cell cycle analysis

The cell cycle was analyzed with a Cell Cycle and Apoptosis Analysis Kit (Beyotime). For cell cycle analysis, samples of 1×10^6^ cells were fixed and permeabilized by the addition of 1 mL of ice-cold 70% ethanol for 15 minutes on ice. After washing, the cells were resuspended in 125 μL of 1.12% (w/v) sodium citrate containing 0.2 mg/mL RNase (Beyotime) and incubated at 37°C for 15 minutes. Next, 125 μL of 1.12% (w/v) sodium citrate containing 50 μg/mL propidium iodide (Beyotime) was added to the cells. Following treatment for 30 minutes at room temperature in the dark, the cells were stored at 4°C until analysis by flow cytometry (FACSCalibur, BD Biosciences). Cell cycle analysis was performed via ModFit LT software.

### Xenograft tumor model and treatment

HCT116 or LS174T cells were inoculated subcutaneously into the flanks of female BALB/c athymic nude mice (2×10^6^ cells per 200 μL or 1×10^7^ per 200 μL). Seven to ten days later, the animals were grouped by randomizing the tumor sizes (approximately 70–120 mm^3^). Five to six animals were used per group. Next, the purified protein or transcription factor decoy DNA plus the Max protein was injected via the tail vein once every 3 days, with 100 μL of total volume per injection. The group of mice receiving physiological saline solution injections was used as a negative control, whereas the group receiving cisplatin injection was used as a positive control. Moreover, the lengths and widths of the tumors were measured every 3 days with a Vernier caliper. The tumor volumes were calculated via the following formula: volume = (length × width^2^)/2 for each animal.

### De novo design of the CEA-binding proteins CEABP1 and CEABP2

The RFdiffusion method was used to construct the scaffolds of the designed CEA-binding proteins, including CEABP1 and CEABP2. By introducing noise into the initially randomly arranged amino acid residues and considering the CEA structure, the process iteratively reduces the noise while simulating protein dynamics in a physiological environment. This results in a scaffold structure that optimally folds and spatially complements the specific binding sites of CEA. During this process, energy functions and geometric constraints are applied to optimize the scaffold, avoiding local potential traps. Each iteration step evaluates and corrects the scaffold on the basis of the structural features of the target protein, making it more compatible with the binding interface or functional regions on the CEA surface. Ultimately, designs with high stability and optimal complementarity are selected.

The ProteinMPNN method was employed to design the amino acid sequences of the designed CEA-binding proteins. During this procedure, the graph neural networks and a message-passing mechanism were utilized to integrate spatial and energy information. This method selects the most appropriate amino acid type for each residue position, ensuring high compatibility between the sequence and scaffold while maintaining excellent folding potential. In the design of CEA-binding proteins, the ProteinMPNN focuses particularly on optimizing the hydrophobic cores, the hydrogen bond networks, and the salt bridges at the protein□protein binding interfaces, significantly increasing the binding affinity and specificity. Through a “scaffold-sequence” two-step approach, a set of candidate CEA-binding proteins with high binding affinity, thermal stability, and expression capacity were designed by inputting parameters such as the three-dimensional structure of the CEA, hotspot residues, and noise reduction scale. The RFdiffusion technology addresses the challenges of spatial geometric constraints and functional residue accommodation, whereas the ProteinMPNN method ensures stable folding of the residue combinations and high-selectivity binding. The synergy between these two technologies enhances the functionality and expressibility of the designed proteins.

Finally, AlphaFold3 and Rosetta were used to exclude poorly folded candidate sequences. Protein□target binding was then evaluated to reduce the false positive rate. AlphaFold3 was used to predict the protein complex structure, focusing on the predicted alignment error (PAE) and root mean square deviation (RMSD) metrics to assess the stability of the interfaces between CEA and the designed CEA-binding proteins. Rosetta was employed to quantify the free energy of the binding interface, analyze the hydrogen bonds and spatial complementarity, and address potential issues through iterative optimization. Using a custom weight function that combines AlphaFold3 evaluation metrics with Rosetta’s energy functions, several sequences were selected through multiple rounds of screening and redesign. Molecular dynamics simulations were performed to observe the dynamic interaction between the designed CEA-binding proteins and the target protein, evaluating the stability of the binding interface and selecting structures with greater dynamic stability and tighter specific binding. Through multiple rounds of evaluation and selection among 10,000 to 100,000 candidate designed sequences in silico, 5 to 10 designed sequences of CEA-binding proteins were selected for experimental validation.

### Molecular dynamics (MD) simulation

AlphaFold3 was used to predict complex structures, and topologies for MD simulations were generated with pdb4amber and tleap, tools from the AMBER suite.

MD simulations were carried out via AMBER, which employs the TIP3P water model and the ff14SB force field for solvent and protein interactions. Two-step energy minimization was performed, first focusing on the solvent and then the whole system. After minimization, the system was gradually heated from 0 K to 300 K over 50 pecoseconds (ps). Equilibration was performed under NPT conditions at 300 K and 1 bar for 50 ns for density stabilization. The equilibrated structures were used for production simulations lasting at least 400 ns to generate structural data. SHAKE was applied after minimization to constrain hydrogen bonds, and PME was used for long-range electrostatics. Trajectory analysis was performed with Cpptraj and MDAnalysis, and visualizations were created via PyMOL.

### Immunofluorescence staining

The cells that were incubated with GFP-tagged CEA-binding proteins (CEABP1 or CEABP2) were fixed with 4% paraformaldehyde at room temperature for 15 minutes. The samples were further incubated with PBS blocking buffer (containing 2% BSA and 2% nonfat milk) at room temperature for 30 minutes. Afterwards, the samples were incubated with primary antibody overnight at 4°C, washed with PBS, and incubated with goat anti-rat-Alexa Fluor 647 (Yeasen) in blocking buffer (1:1000 dilution) at room temperature for 60 minutes. The stained samples were washed with PBS, and the nuclei were stained with Hoechst 33342 (MA0126, Meilunbio, Dalian, China; 1:100 dilution in PBS). The samples were mounted on slides with Fluoromount-G™ mounting medium (Yeason).

### Hematoxylin & eosin (H&E) staining

The paraffin-embedded slides were first deparaffinized with xylene, followed by a series of grades of ethanol, and finally with water. The tissues on the slides were then incubated with a hematoxylin solution for one minute and washed with tap water until the water was clear. Next, the tissues were counterstained in an eosin solution for 15□seconds. After counterstaining, the tissue slides were immediately transferred to 95% ethanol and further dehydrated with 100% ethanol and xylene. Finally, the slides were mounted with neutral balsam mounting medium and air-dried before imaging.

### Antibodies

The antibody for CEA (Proteintech Cat#: 10421-1-AP) and Alexa Fluor 594-conjugated goat anti-rabbit IgG (H+L) (ABclonal Cat#: AS039) were used in this study.

### Statistical analysis

All the graphs were prepared via GraphPad Prism 9 software, and statistical analysis was also carried out via GraphPad Prism 9 software to perform one-way ANOVA, two-sided t test, or two-way ANOVA. The error bars indicate the standard error of the mean (SEM). A p value < 0.05 is considered statistically significant, where all statistically significant values shown in the figures are indicated as follows: * p < 0.05, ** p < 0.01, *** p < 0.001, and **** p < 0.0001.

## Supporting information

Supplementary Videa

## Acknowledgements

This research was funded by the National Key R&D Program of China (grant numbers 2020YFA0907300 and 2022YFA0912201) and the National Natural Science Foundation of China (grant number 32170030).

## Author contributions

G.W. conceived the project. W.W. designed, performed, and analyzed the protein purification, cell assays, and xenograft tumor mouse model experiments. X.S. performed the protein design, protein structure prediction, and molecular dynamics simulation. G.W., W.W., and X.S. wrote the paper.

