## Supplementary Figures for "Targeted inhibition of colorectal carcinoma using a designed CEA-binding protein to deliver TCF/LEF transcription factor decoy DNA and p53 protein"

**SUPPLEMENTARY INFORMATION**

**SUPPLEMENTARY FIGURES**

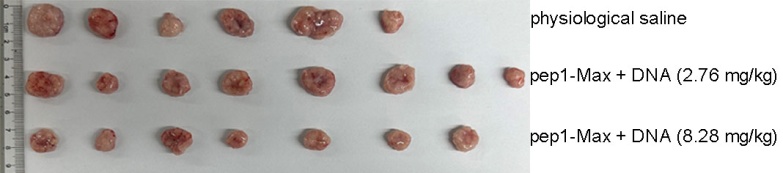

**Figure S1 Representative images of HCT116 cell xenograft tumors at the experimental endpoint corresponding to Figure 1F.**

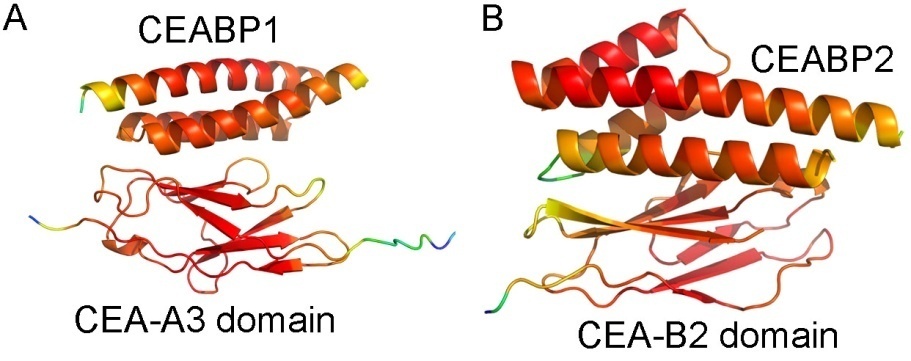

**Figure S2 Confidence levels of the predicted structure of CEABP1 in complex with the CEA-A3 domain and that of CEABP2 in complex with the CEA-B2 domain were very high.**

(A) The predicted local distance difference test (pLDDT) scores of CEABP1 and the A3 domain of CEA were mapped to the predicted structure of their complex. Higher confidence values are shown in warmer colors.
(B) The pLDDT scores of CEABP1 and the B2 domain of CEA were mapped to the predicted structure of their complex. Higher confidence values are shown in warmer colors.

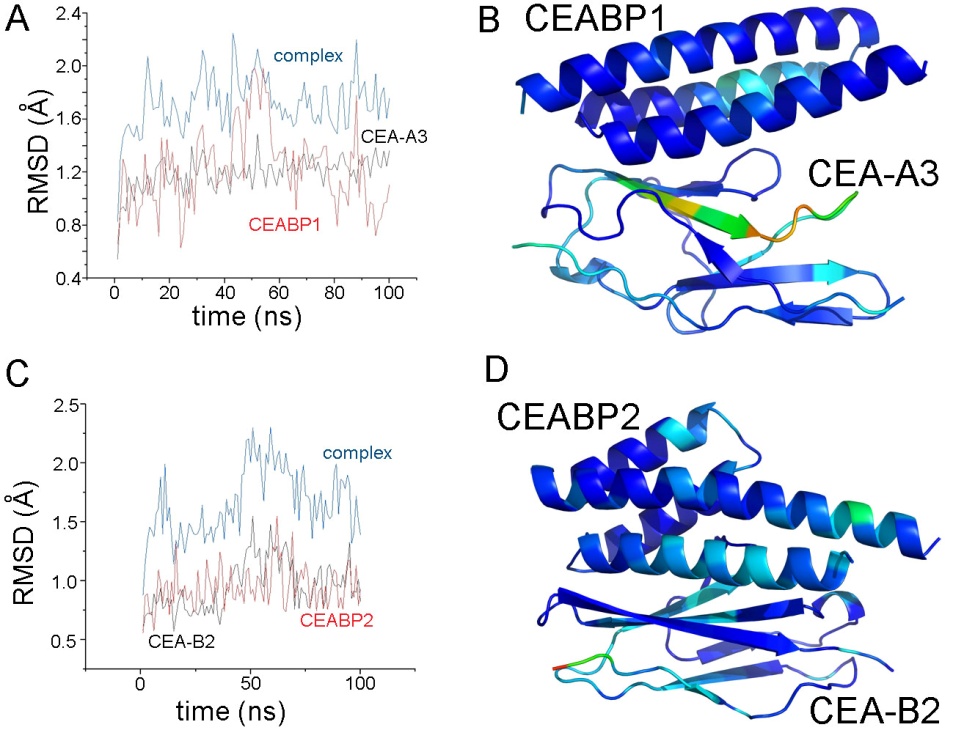

**Figure S3. Molecular dynamics simulation results showing that the complex between CEABP1 and the CEA-A3 domain and that between CEABP2 and the CEA-B2 domain were stable.**

(A) Time-dependent root mean square deviation (RMSD) of the complex between CEABP1 and the CEA-A3 domain from molecular dynamics simulation.
(B) The structure of CEABP1 in complex with the CEA-A3 domain is colored according to the root mean square fluctuation (RMSF) values from the molecular dynamics simulation. Greater fluctuations are shown in warmer colors.
(C) RMSD of the complex between CEABP2 and the CEA-B2 domain from molecular dynamics simulation.
(D) The structure of CEABP2 in complex with the CEA-B2 domain is colored according to the RMSF values from the molecular dynamics simulation. Greater fluctuations are shown in warmer colors.

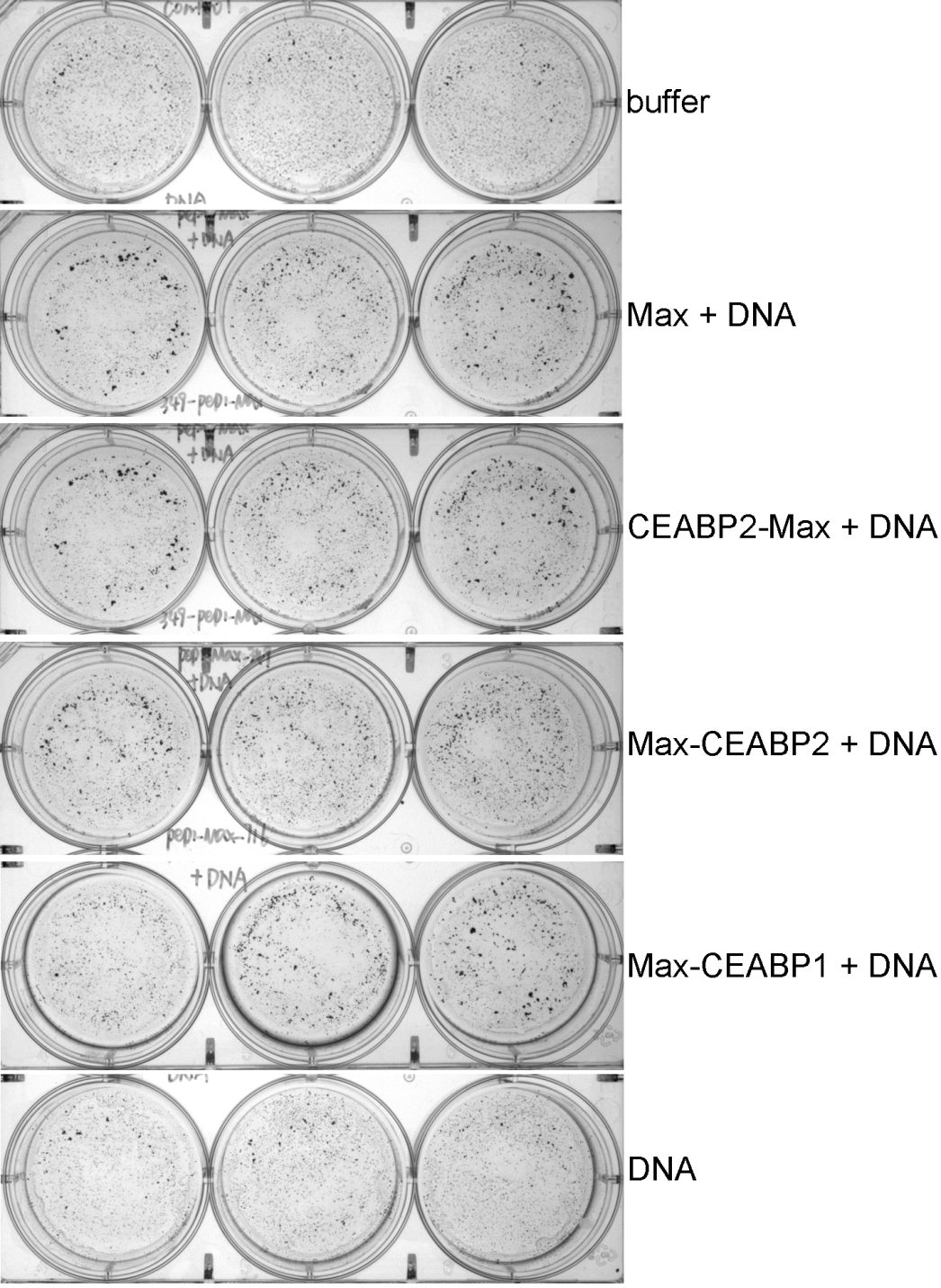

**Figure S4. Colony formation assay results showing that fusing CEABP1 to pep1-Max caused the delivered TCF/LEF TFD DNA to inhibit LS174T cell growth more strongly than it did in the absence of CEABP1, corresponding to Figure 3H.**

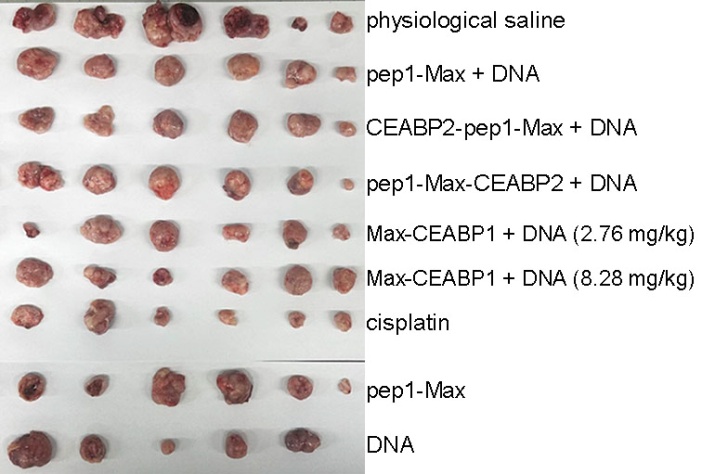

**Figure S5. Representative images of LS174T tumors at the endpoint, corresponding to Figure 3I.**

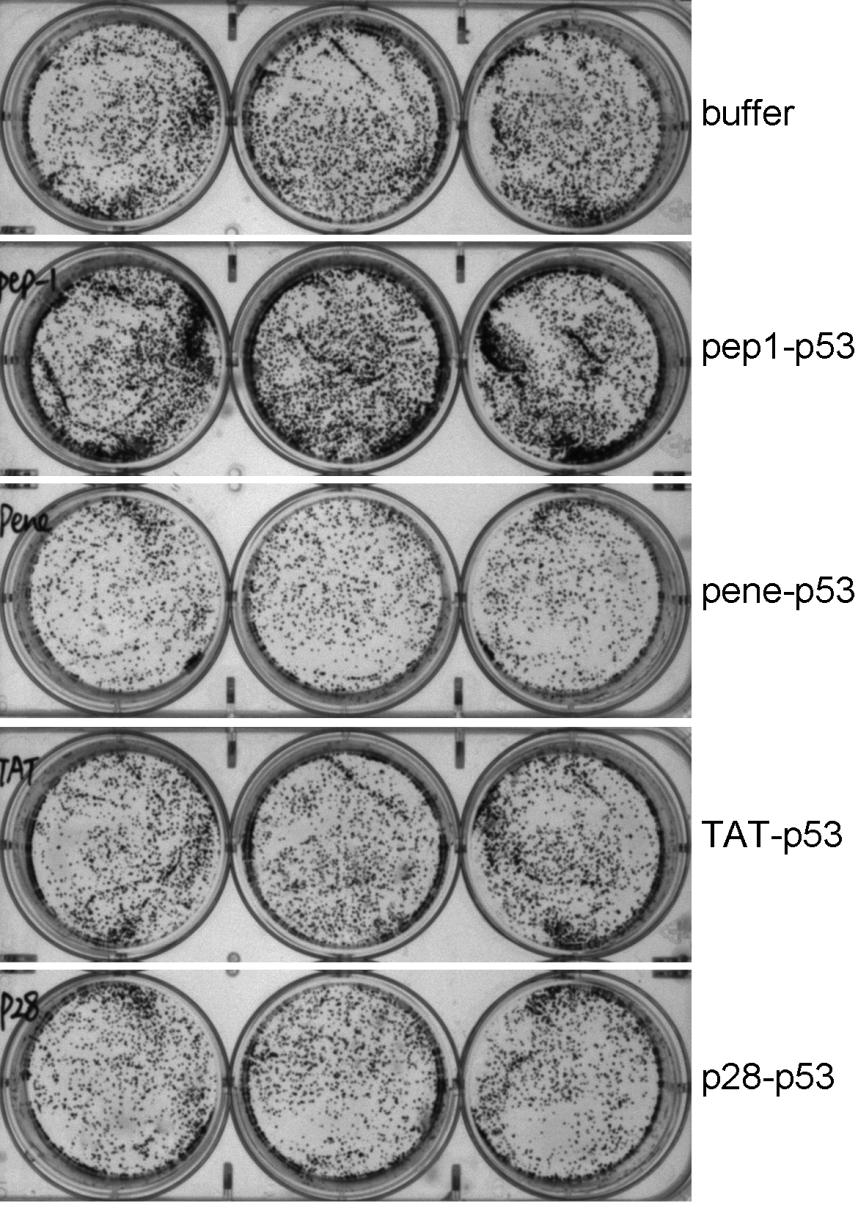

**Figure S6. p28-p53 exhibited the strongest inhibition of cell proliferation when delivered into HCT116 cells, as examined by the colony formation assay, the statistics of which is shown in Figure 4B.**

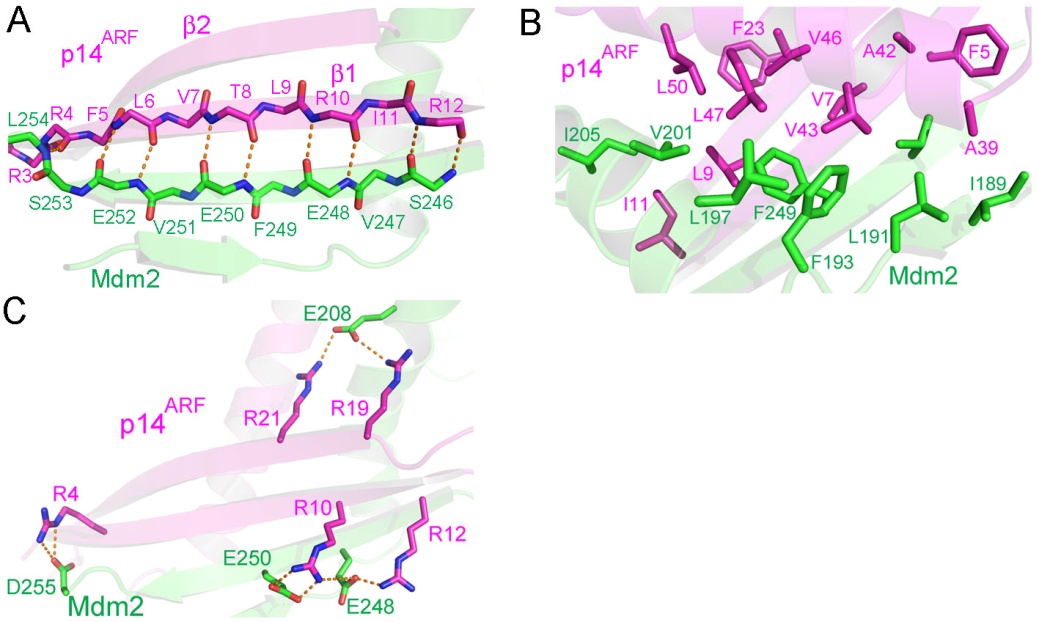

**Figure S7. Predicted interactions between p14^ARF^ and Mdm2.**

(A) Main-chain hydrogen bonding interactions between p14^ARF^ and Mdm2.
(B) Hydrophobic interactions between p14^ARF^ and Mdm2.
(C) Electrostatic interactions between p14^ARF^ and Mdm2.

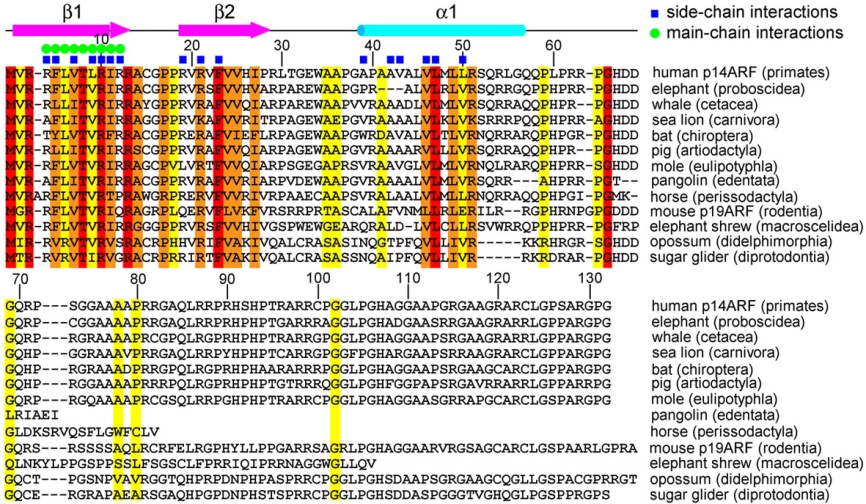

**Figure S8. N-terminal 63 residues of p14^ARF^ are highly conserved.** Multiple sequence alignments were performed for representative p14^ARF^ orthologs from humans, African savanna elephant (*Loxodonta africana*, accession number: XP_064146496), gray whale (*Eschrichtius robustus*, XP_068408523), Steller sea lion (*Eumetopias jubatus*, XP_027970991), bat (*Myotis myotis*, KAF6314295), pig (*Sus scrofa*, CAD53376), Iberian mole (*Talpa occidentalis*, XP_054545170), Chinese pangolin (*Manis pentadactyla*, KAI5158232), Przewalski’s horse (*Equus przewalskii*, XP_008508663), mouse p19^ARF^, Cape elephant shrew (*Elephantulus edwardii*, XP_006881402), gray short-tailed opossum (*Monodelphis domestica*, NP_001028145), and sugar glider (*Petaurus breviceps papuanus*, XP_068943629).

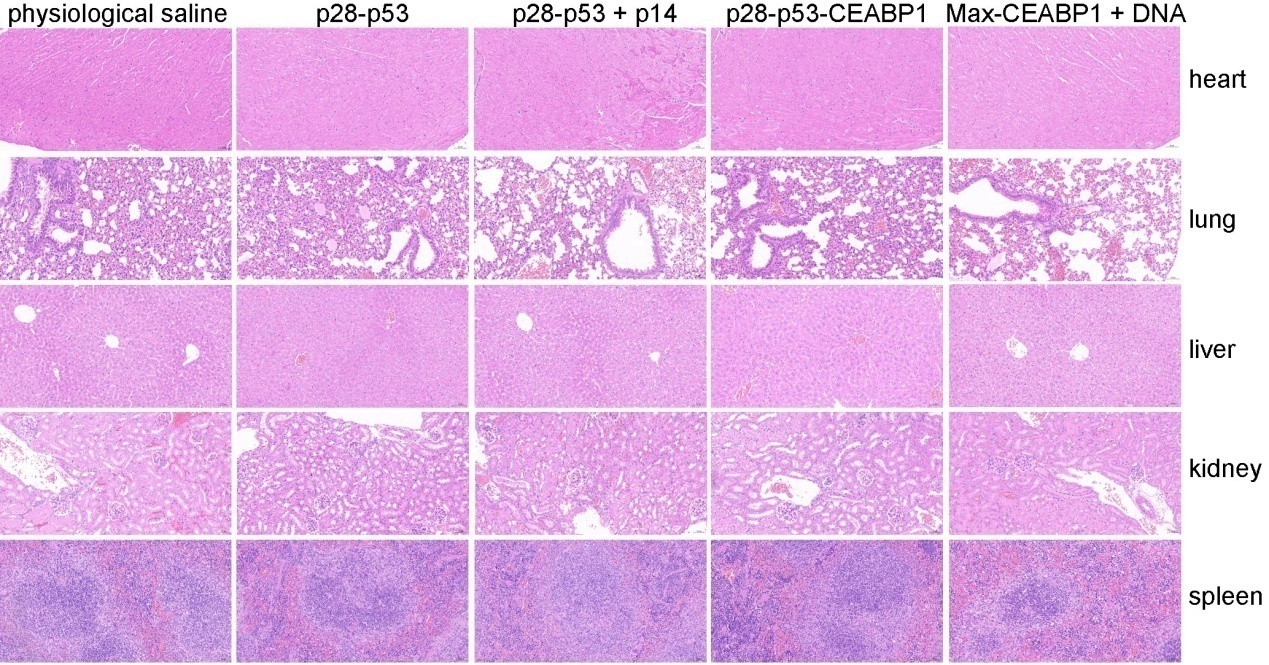

**Figure S9. Hematoxylin and eosin (H&E) staining analysis of harvested major organs (heart, liver, spleen, lung, and kidney) revealed no detectable histopathological abnormalities in mice treated with p28-p53 or p28-p53 together with p28-p14^ARF^, p28-p53-CEABP1, or pep1-Max-CEABP1 together with TCF/LEF TFD DNA.**

**SUPPLEMENTARY MOVIE**

**Movie S1. The predicted structure of p14^ARF^ (residues 1-63) in complex with human Mdm2 (residues 186-255) was subjected to molecular dynamics simulation, and the complex between p14^ARF^ (1-63) and Mdm2 (186-255) was stable during the molecular dynamics simulation process.**
